# A mutation in vesicular acetylcholine transporter increases tubulin acetylation compromising synaptic vesicle transport

**DOI:** 10.1101/2024.06.06.597842

**Authors:** Cheng-Shan Kuo, Meng-Chieh Wang, Vignesh M. Ruckmani, Muhammad S. Khawaja, Odvogmed Bayansan, Syed Nooruzuha Barmaver, Prerana Bhan, Oliver Ingvar Wagner

## Abstract

Kinesin-3 UNC-104(KIF1A) is the major anterograde axonal transporter of synaptic vesicles and is expressed pan-neuronally. Genetic defects in this molecular motor are linked to KIF1A-associated neurological disorders (KAND) encompassing Charcot-Marie-Tooth (CMT) disease and hereditary spastic paraplegia (HSP). From a candidate screen for genes causing neurotransmission defects in *C. elegans* and simultaneously affecting post-translational modification of tubulin, we identified allele *unc-17*(*e245*) significantly elevating tubulin acetylation *in vitro* and *in vivo*. UNC-17 encodes for a VAChT (vesicle acetylcholine transporter) and its human ortholog SLC18A3 is implicated in Alzheimer’s and Huntington’s disease. To exclude secondary effects of the *unc-17* mutation, we tracked UNC-104 and RAB-3 motility in the non-cholinergic ALM neuron. With upregulated tubulin acetylation in ALM (anterior lateral microtubule) neurons in *unc-17*(*e245*) strains (visualized by immunostaining), motility of both, motor and its cargo, is significantly compromised. However, motility of UNC-104 improves when knocking down alpha-tubulin acetyltransferase MEC-17(ATAT1) in *unc-17*(*e245*) strains and, conversely, is negatively affected when overexpressing MEC-17 in wild type animals. UNC-17 and UNC-104 are co-expressed and colocalize in cholinergic head neurons, suggesting a functional motor-cargo relationship. Strikingly, *mec-17* knockdown significantly enhances their colocalization, while *unc-17* knockdown reduces UNC-104/MEC-17 colocalization in ALM neurons. Direct interactions were confirmed by bimolecular fluorescence complementation (BiFC) and co-immunoprecipitation (Co-IP): MEC-17 depletion strengthens UNC-104/UNC-17 association and stabilizes the motor-cargo complex, whereas *unc-17* knockdown attenuates UNC-104/MEC-17 interactions, as evidenced by diminished Co-IP signals. We propose that *unc-17* knock-down releases MEC-17 from the transport complex, thereby enhancing its tubulin acetyltransferase activity. This elevated acetylation may disrupt motor motility, ultimately impairing synaptic vesicle trafficking. This dynamic interplay, modulated by tubulin acetylation, highlights a regulatory axis involving UNC-104, UNC-17, and MEC-17 across cholinergic and glutamatergic neurons, offering new insights into axonal transport defects relevant to KIF1A-associated neurological disorder (KAND).

## INTRODUCTION

Neurons are highly polarized cells with long projections (termed axon and dendrite) radiating from their cell bodies (also called soma). Since most proteins are synthesized in the soma, a proper intra-cellular transport system must be maintained to ensure that proteins can reach the periphery of neurons for several critical functions such as synaptogenesis, synaptic transmission, axon growth and pathfinding (Guillaud et al., 2020). Axonal transport is a bi-directional transport along microtubule (MT) tracks in which molecular motors of the kinesin class transport cargos to distal regions of neurons (anterograde) or via dyneins back to the soma (retrograde) (Hirokawa et al., 2010). Impaired axonal transport may lead to the accumulation of “disease proteins” such as tau, neurofilaments, or alpha-synuclein all linked to neurological diseases such as Huntington’s disease (HD), Alzheimer’s disease (AD) and Parkinson’s diseases (PD) (Millecamps & Julien, 2013). Kinesin-3 UNC-104(KIF1A) is the major axonal transporter of RAB-3, synaptotagmin, synaptobrevin and synaptophysin (Hall & Hedgecock, 1991; Maeder et al., 2014), and genetic defects in this motor are linked to KIF1A-associated neurological disorders (KAND) encompassing Charcot-Marie-Tooth (CMT) disease and hereditary spastic paraplegia (HSP) (Chiba et al., 2023; Xu et al., 2018). UNC-104 regulates dendrite branch morphogenesis (Kern et al., 2013; Zhang et al., 2016), density of synapses (Niwa et al., 2016) and is involved in synaptic aging and memory (Li et al., 2016). Homologies between human KIF1A and *C. elegans* UNC-104 are high with 72% positives and 56% identities. UNC-17 encodes for a VAChT (vesicle acetylcholine transporter) that associates with synaptic vesicles (Alfonso et al., 1993) and its human ortholog SLC18A3 (solute carrier family 18 member A3) is linked to AD, HD, pulmonary inflammation and congenital myasthenic syndrome 21 (Banzato et al., 2021; Ferreira-Vieira et al., 2016; Lamond et al., 2021; Smith et al., 2006; Suzuki et al., 2001; Zadrozny et al., 2024). Homologies between human VAChT (SLC18A3) and *C. elegans* UNC-17 are 47% positives and 61% identities. GTP-bound Rab3 acts as an adaptor of UNC-104 to allow the transport of various vesicular cargos. At the nerve terminal, Rab3 regulates synaptic vesicle docking and fusion as well as facilitates neurotransmitter release (Binotti et al., 2016). Homologies between human Rab3A and *C. elegans* RAB-3 are high with 81% positives and 73% identities.

Tubulin is the basic component of MTs and their C-terminal ends form unstructured tails emanating from the MTs. Various enzymes can add or remove specific amino acids such as glutamate and tyrosine to and from these tails leading to post-translational modifications (PTMs) of tubulin (Janke & Magiera, 2020). PTMs regulate MT stability and the attachment/detachment rates of MAPs (microtubule-associated proteins) and molecular motors (Genova et al., 2023). Another PTM is acetylation that only occurs at lysine-40 of α-tubulin inside the lumen of MTs (Li & Yang, 2015). Acetylation of tubulin weakens the interaction between protofilaments that increases MT flexibility protecting MTs from breakage under stress (Eshun-Wilson et al., 2019). Acetylation of tubulin also increases kinesin-1 attachment rates, therefore promoting axonal transport (V. K. Godena et al., 2014; Reed et al., 2006). However, the relationship between tubulin acetylation and kinesin-1 motility is still debated, with some studies showing no direct effect of acetylation on kinesin-1 velocity or run length (Kaul et al., 2014; Walter et al., 2012). Acetylation of tubulin is catalyzed by α-tubulin acetyltransferase (ATAT1) (Janke & Magiera, 2020) which is MEC-17 in *C. elegans* (49% positives/64% identities). A mutation in MEC-17 in *C. elegans* leads to variable numbers of protofilaments in touch receptor neurons (TRN) (Cueva et al., 2012). Besides, MEC-17 regulates the development of synaptic branches by its non-enzymatic domain, and in *mec-17* mutants, synaptic vesicles are decreased at synapses in TRN (Teoh et al., 2022). Notably, reduced levels of tubulin acetylation are found in PD, HD and CMT and all of these diseases are linked to disturbed axonal transport. Particularly, decreased tubulin acetylation levels are detected in HD mouse models and human patient samples. However, the relationship between tubulin acetylation and axonal transport defects in HD is complex, such as restoring acetylation levels alone does not fully rescue transport impairments (d’Ydewalle et al., 2011; Dompierre et al., 2007; V. K. Godena et al., 2014). While some studies reveal decreased tubulin acetylation levels in PD mouse models and human patient samples, others have found no significant changes or even increased acetylation levels (Mazzetti et al., 2024).

The direct relation between synaptic transmission and axonal transport is evident from various studies. For example, in *C. elegans*, kinesin-1 UNC-116(KIF5) regulates the delivery and removal of AMPA receptor GLR-1 at postsynapses. Mutated UNC-116 causes changes in the surface expression of GLR-1 directly modulating electrophysiological signals (Fan & Lai, 2022; Hoerndli et al., 2013). Conversely, a relation between PTM and axonal transport (and in turn synaptic transmission) has been described, such as that the reduction of tubulin polyglutamylation alters the neuronal distribution of KIF1A which in turn directly modulates synaptic transmission (Ikegami et al., 2007). Likewise, activation of AMPAR leads to increased levels of polyglutamylation directly affecting KIF5-based transport (Maas et al., 2009).

In this study, we carried out a candidate screen of various *C. elegans* strains known to display synaptic transmission defects and monitored changes in tubulin PTM in these mutants. We then investigated changes in transport of synaptic vesicles in neurons with elevated tubulin acetylation to provide a link between synaptic transmission, tubulin PTM and synaptic vesicle transport.

## RESULTS

### Synaptic transmission defects in cholinergic neurons increases tubulin acetylation

Tubulin PTMs are crucial for regulating microtubule dynamics that play a vital role in neuronal health and synaptic function. In a candidate screen for *C. elegans* mutant strains that exhibit synaptic transmission defects and at the same time causing changes in tubulin PTM, we employed wild type N2 worms as well as 7 different mutant strains: *dat-1* with defects in dopamine neurotransmission (Nass et al., 2002), *eat-4* with defects in glutamate neurotransmission (Avery, 1993), *unc-17* with defects in ACh neurotransmission (Sandoval et al., 2006), *unc-25* with defects in GABA neurotransmission (McIntire et al., 1993), *tbh-1* with defects in octopamine neurotransmission (Alkema et al., 2005), *tdc-1* with defects in tyramine neurotransmission (Alkema et al., 2005), and *tph-1* with defects in serotonin neurotransmission (Sze et al., 2000). Western blots from whole worm lysates revealed a 1.54-fold upregulation in tubulin acetylation in the *unc-17* strain whereas changes in other strains remained insignificant (Fig. 1A+B, Suppl. Fig. S3A). Interestingly, tubulin tyrosination remained unchanged in all strains (Fig. 1A, Suppl. Figs. S1+S3B). We did not screen for changes in glutamylation since in *C. elegans* this type of PTM is mostly seen in cilia or male sensory neurons (O’Hagan et al., 2022). Employing whole-mount immunostaining, we then demonstrate that acetylation is 3-fold increased in ALM neurons of *unc-17*(*e245*) mutants (Fig. 1C-E). The gene *unc-17* encodes for a synaptic vesicular acetylcholine transporter (VAChT) and the hypomorphic allele *e245* carries a G/C substitution in the 3rd (and last) exon of *unc-17* causing the worms to coil-up at all developmental stages with impeded locomotion associated to neurotransmission defects (Alfonso et al., 1993; Sandoval et al., 2006; Zhu et al., 2001). While direct stability assays for e245 are unavailable, Alfonso et al., 2013 noted that other unc-17 mutations (md1447, p279) reduce UNC-17 immunostaining, suggesting potential protein instability or degradation. Immunostaining of *unc-17*(*e245*) worms with a monoclonal anti-UNC-17 antibody shows reduced expression of UNC-17 in various cholinergic neurons (Mathews et al., 2021; Sandoval et al., 2006).

**Figure 1:**
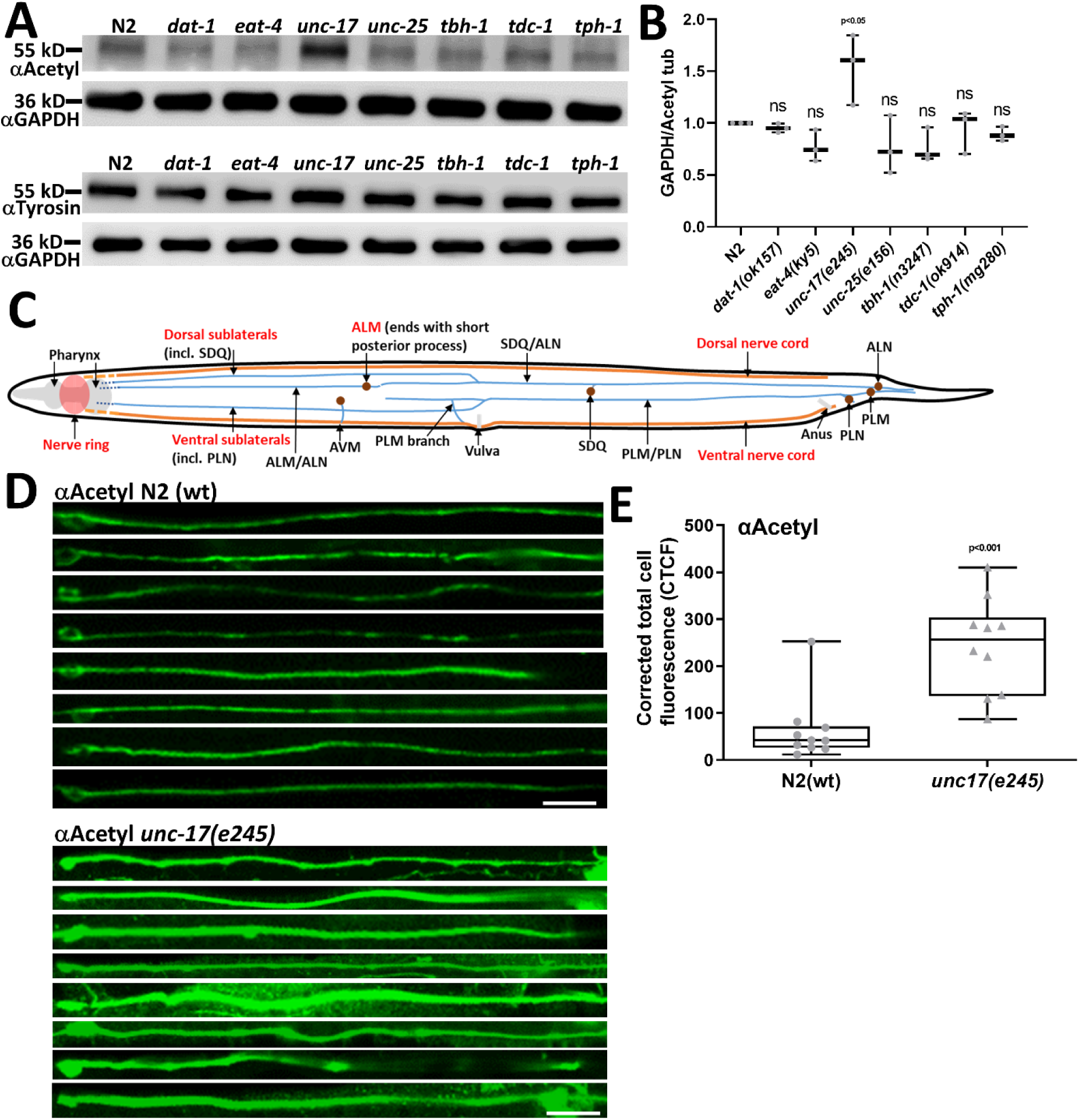
Candidate screen for genes affecting neurotransmission and simultaneously tubulin PTM in *C. elegans*. (A) Western blot analysis to investigate changes in tubulin acetylation (upper panel) and tyrosination (lower panel) in various *C. elegans* strains carrying mutations reported to cause synaptic transmission defects. (B) Quantification of band signal intensity from (A, upper panel) using ImageJ software. For tyrosination quantification (A, lower panel) refer to Suppl. Fig. S1. Western blots repeated in triplicates. (C) Simplified scheme of the nervous system in *C. elegans*. (D) Whole mount immunostaining using an anti-acetylated tubulin antibody. The panels display representative confocal images of stacked ALM neurons (C). Upper panel: ALM neurons from N2 wild type worms. Lower panel: ALM neurons from worms with mutations in the *unc-17* gene (*e245* allele). (E) Quantification of images shown in (D) using Image J. Total analyzed neurons per strain: 10. Oneway ANOVA with Dunnett’s test for multiple comparisons utilized in (B). Two-tailed, unpaired Student’s t-test in (E). Box and whisker plots represent maximum value, upper quartile, median, lower quartile and minimum value. Scale bars: 10 µm.

### Increased tubulin acetylation negatively affects motility of UNC-104 and its cargo RAB-3

To understand the potential impact of altered tubulin PTM on axonal transport, we analyzed the motility of the major axonal transporter of synaptic vesicles UNC-104 and its cargo RAB-3 in neurons of living *C. elegans* animals. To exclude secondary effects of the synaptic transmission defect in *unc-17* (likely to affect transport), we use the glutamatergic ALM neuron for our observation. The ALM neuron is one of several TRN possessing a particularly long axon (extending up to 0.5 mm), and that can be easily identified among the other neurons in nematodes (Fig. 1C). Note that in previous studies, we have shown that the characteristics of UNC-104 motility are comparable among others TRN such as PLM neuron as well as neurons from the sublateral system (Tien et al., 2011; Wagner et al., 2009; Wu et al., 2016). Intriguingly, motility of both motor (*e1265*;*Punc-104::unc-104::mrfp*) and cargo (N2;*Punc-104::mCherry::RAB-3*) was largely compromised by the *unc-17* mutation (Fig. 2). Both anterograde and retrograde velocities were reduced (Fig. 2A) reflected by an increase in pausing (Fig. 2B). Also, single run length events (Fig. 2C), as well as total run length (Fig. 2D) was negatively affected by the *unc-17* mutation in both directions (see also Fig. 2E). Since UNC-104 is an anterograde motor, the reduction in retrograde directions is likely based on dynein motility. The “paradox of co-dependence model” explains well this finding: motors are mutually activated by mechanically pulling on each other in opposing directions. If this mechanical activation is lost or reduced, motor activation is diminished (Hancock, 2014). We have seen and reported this phenomenon on various other publications (Vinay K. Godena et al., 2014; Muniesh et al., 2020; Tien et al., 2011; Wu et al., 2016).

**Figure 2:**
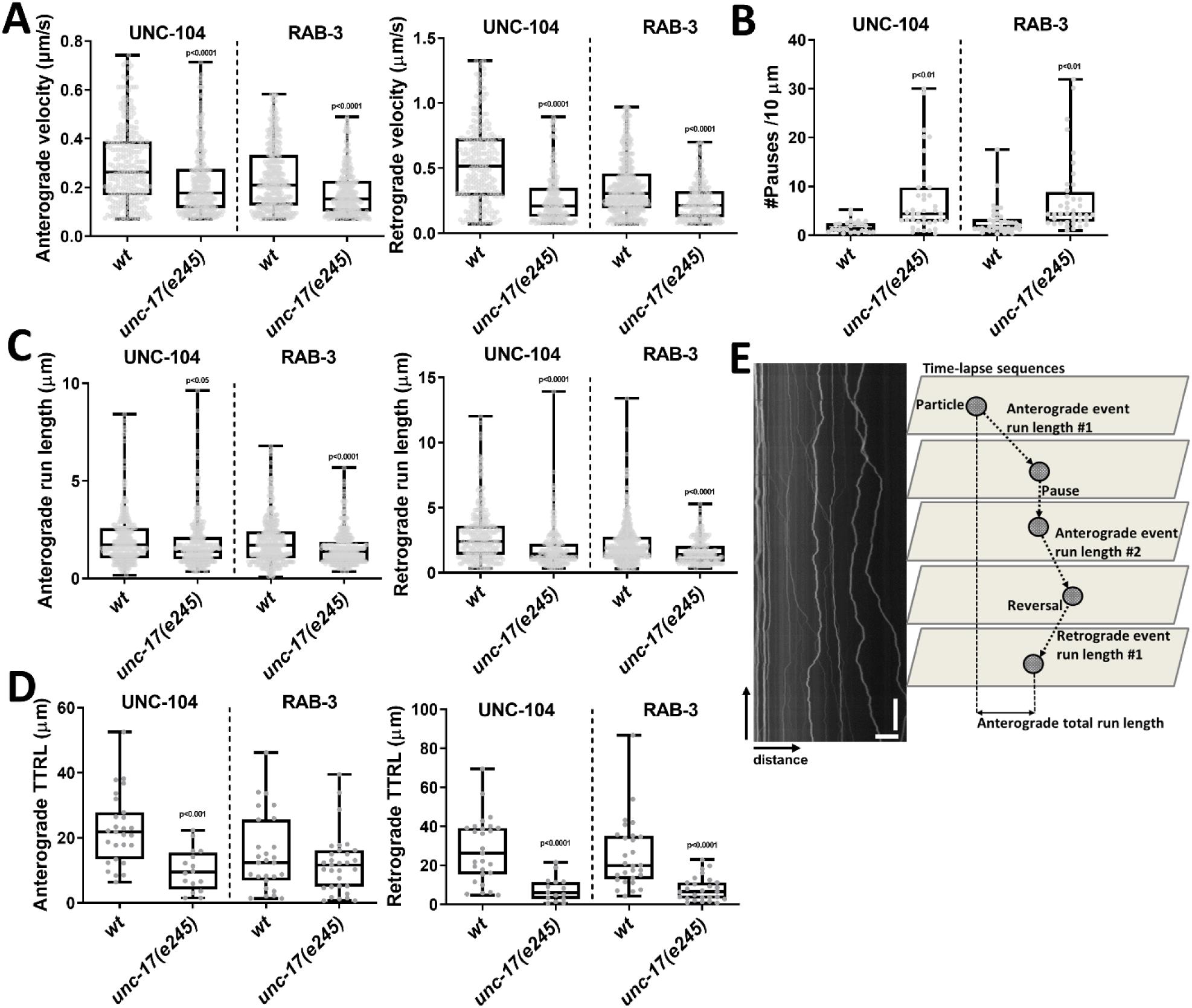
Motor and cargo motility analysis in neurons of living *C. elegans* animals depending on an *unc-17* mutation. (A) Anterograde and retrograde velocities of kinesin-3 UNC-104 (UNC-104::mRFP) and its major cargo RAB-3 (mCherry::RAB-3) in either wild type N2 or *unc-17*(*e245*) mutants. (B) Number of pauses per 10 µm. (C) Anterograde and retrograde event run lengths. (D) Anterograde and retrograde total run lengths (TTRL). Analyzed events: UNC-104(wt) = 1106, UNC-104(*unc-17*) = 938, RAB-3(wt) = 783 and RAB-3(*unc-17*) = 933. Biological replicates: 20 worms per condition. Total vesicles analyzed: UNC-104(wt) = 27, UNC-104(*unc-17*(*e245*)) = 29, RAB-3(wt) = 34, RAB-3 (*unc-17*(*e245*)) = 31. (E) (Left) Kymograph and (right) graphical depiction of kymograph analysis with various motility parameters. Two-tailed, unpaired Student’s t-test with *p<0.05, **p<0.01, ***p<0.001, ****p<0.0001. Box and whisker plots represent maximum value, upper quartile, median, lower quartile and minimum value. Scale bars: vertical = 30 s, horizontal = 10 µm.

In previous studies, we identified that UNC-104 tends to form dynamic aggregates along neurons which likely station for motor/cargo exchange (Tien et al., 2011; Wagner et al., 2009). We found that in *unc-17*(*e245*) motor and cargo aggregates are significantly reduced and appear more diffuse (Fig. 3B+C, E-J). Also, both motor and cargo travel significantly shorter distances in neurons (Fig. 3A+D), an effect that can be visibly seen in neurons (Fig. 3B+C).

**Figure 3:**
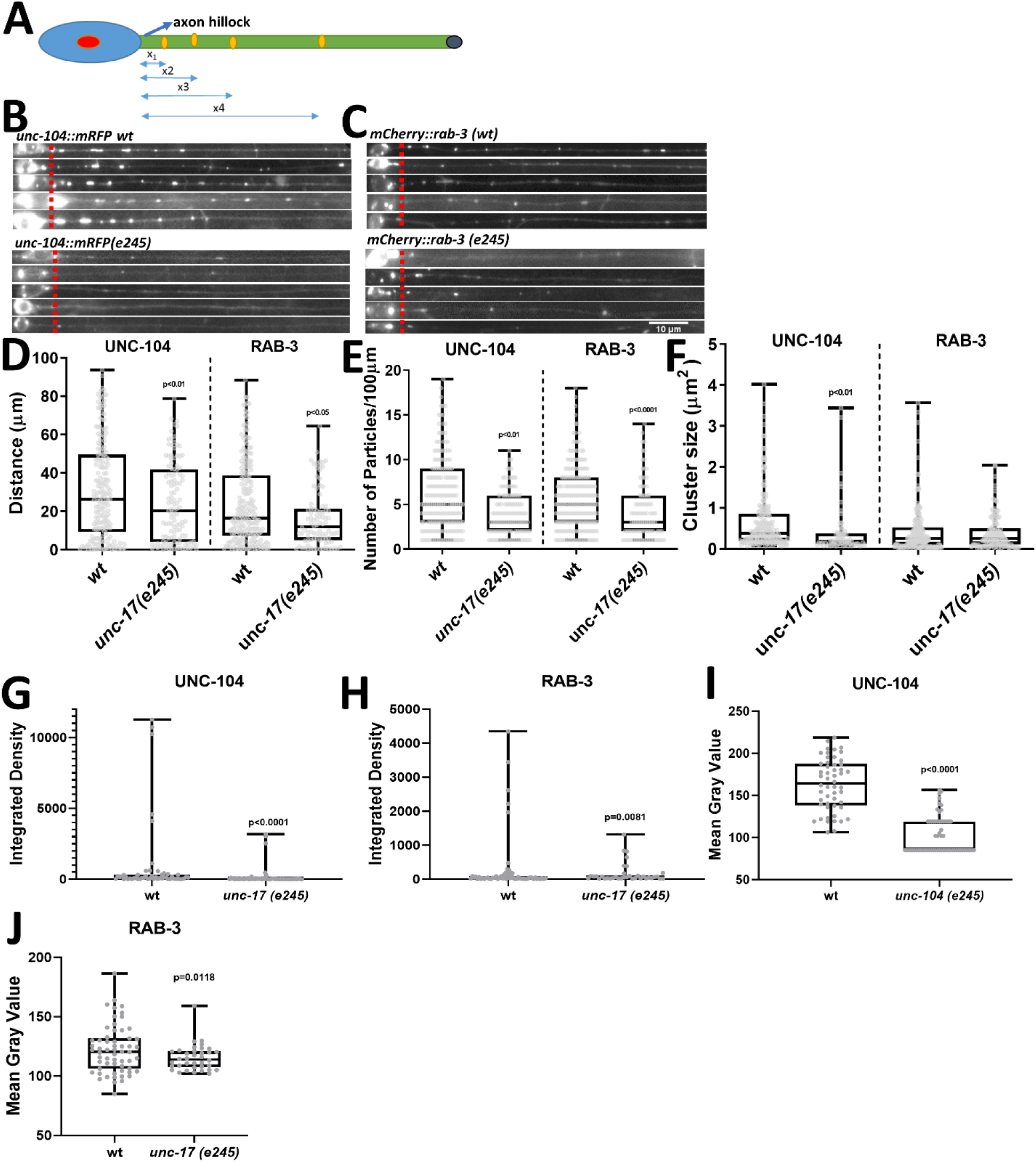
Changes in accumulation propensities and distribution patterns of motor and cargo in neurons caused by the *unc-17* mutation. (A) Schematic diagram of a neuron with blue double-sided arrows indicating the distances traveled from the axon hillock to farther distal regions. (B) Fluorescence images of digitally straightened and stacked ALM neurons taken from different worms expressing UNC-104::mRFP in either wild type or *unc-17* mutants. (C) Representative fluorescence images of stacked ALM neurons taken from different worms expressing mCherry::RAB-3 in either wild type or *unc-17* mutants. (D) Quantification of traveled distances of motor and cargo. (E) Quantification of motor and cargo densities (# particles per 100 µm). (F) Quantification of motor and cargo particles size. (G-J) Quantification of morphology of motor and cargo particles. respectively. Total analyzed neurons per strain: 20. Two-tailed, unpaired Student’s t-test. Box and whisker plots represent maximum value, upper quartile, median, lower quartile and minimum value. Scale bar: 10 µm.

### MEC-17 knockdown rescues the negative impact of the unc-17 mutation *in vivo* and *in vitro*

We then attempted to reduce acetylation in TRNs of *unc-17*(*e245*) worms by knocking down MEC-17, the acetylase responsible for tubulin acetylation in *C. elegans* TRNs (Cueva et al., 2012). As expected, mec-17 knockdown (KD) largely reversed the negative effects on UNC-104 motility in ALM of *unc-17* mutants: velocity significantly increased in both directions (Fig. 4A+B), pausing significantly decreased (Fig. 4C) and anterograde total run length significantly increased (Fig. 4D). Conversely, overexpressing MEC-17 in wild type animals led to comparable effects as measured in *unc-17* mutants (Fig. 4, wt;MEC-17 *OEX*). Western blots confirm the reduction of tubulin acetylation in the *unc-17*(*e245*);*mec-17 KD* strain and the increase in tubulin acetylation in the wt;MEC-17 *OEX* strain (Fig. 4G+H, Suppl. Fig. S3C). Note that we were unable to rescue *e245* effects by overexpressing deacetylase HDA-6 (Suppl. Fig. S4A-D) which is a homolog of human deacetylase HDAC6 (human histone deacetylase 6) known to reverse the action of ATAT1 (Janke & Magiera, 2020). Nevertheless, we also inhibited ACh transmission in wild type animals using the drug levamisole (an AChR agonist) revealing comparable effects on motor motility and tubulin acetylation as determined for the *unc-17* mutation alone (Fig. 4+Suppl. Fig. S5, wt;Levamisole 1mM). With these data, we are first to disclose the direct negative effect of tubulin acetylation on UNC-104(KIF1A)-mediated axonal transport.

**Figure 4:**
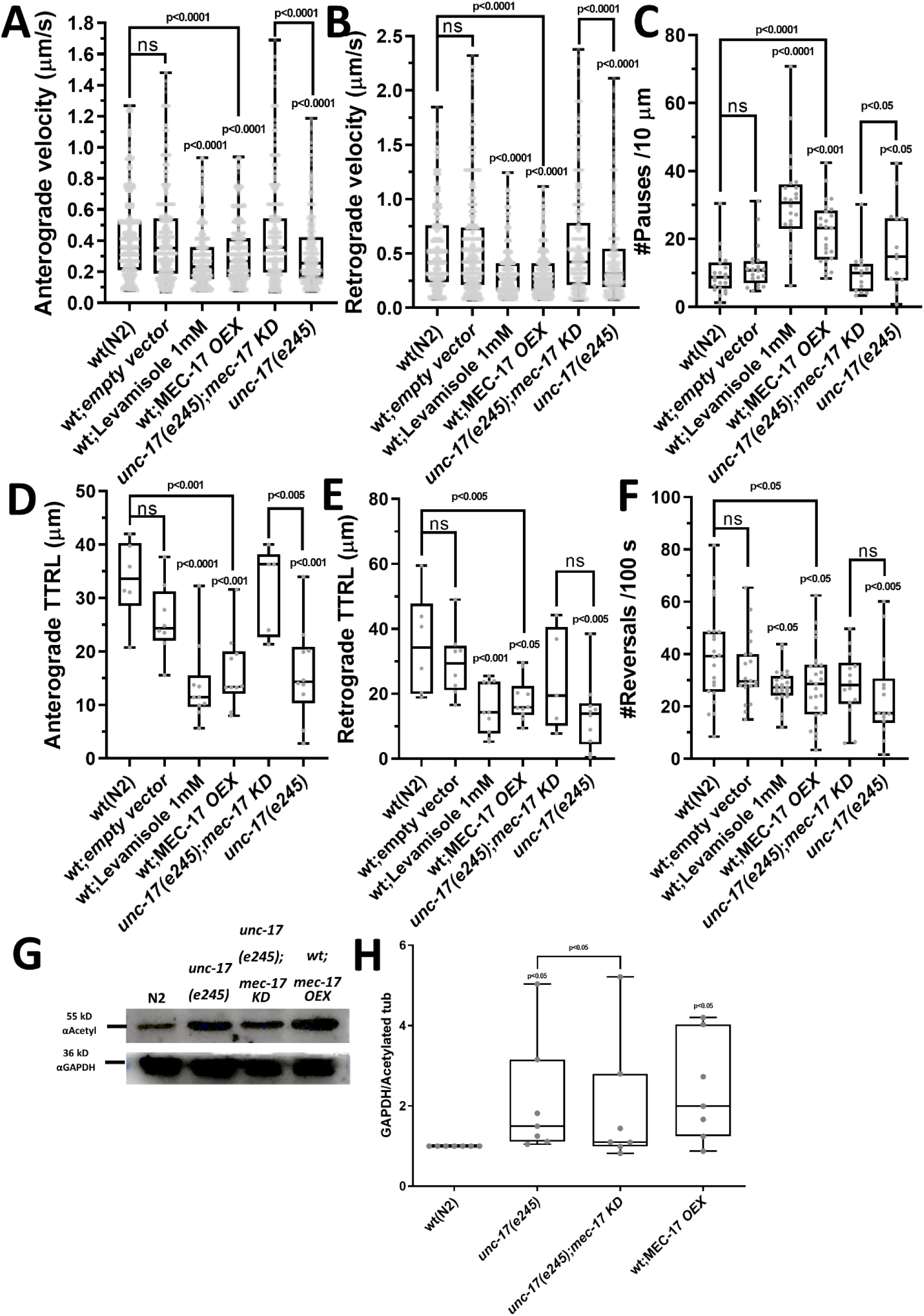
Motor motility after overexpressing/knockdown of acetyltransferase or after adding an ACh neurotransmission inhibitor. (A-F) Quantification of various motility parameters of UNC-104::mRFP in neurons of living *C. elegans* animals. ACh neurotransmission inhibitor levamisole was added at 1mM to NGM agar and acetyltransferase MEC-17 was either overexpressed (OEX) or knocked down (KD). (A) Anterograde and (B) retrograde velocity. (C) Number of pauses per 10 µm. (D) Total anterograde and (E) retrograde run lengths. (F) Quantification of directional changes (motor reversals, Fig. 2E). (G) Western blot exposing the *mec-17* KD and OEX effect on tubulin acetylation. (H) Band density quantification of gel shown in (G). Analyzed events (n): (A+B) wt(N2) = 2270, wt;EV = 2092, wt;Leva = 2047, wt;MEC-17 *OEX* = 2115, *unc-17*(*e245*);mec-17 *KD* = 1389, *unc-17*(*e245*) = 1093. (C+F) wt(N2) = 44, wt;EV = 40, wt;Leva = 48, wt;MEC-17 *OEX* = 50, *unc-17*(*e245*);mec-17 *KD* = 28, *unc-17*(*e245*) = 32. (D+E) wt(N2) = 12, wt;EV = 16, wt;Leva = 20, wt;MEC-17 *OEX* = 20, *unc-17*(*e245*);mec-17 *KD* = 10, *unc-17*(*e245*) = 20. Biological replicates: 20 worms per condition. Total vesicles analyzed: UNC-104(wt;MEC-17 OEX) = 101, UNC-104(wt;empty vector) = 105, UNC-104(*unc-17*(*e245*)) = 62, UNC-104 (*unc-17*(*e245*);mec-17 KD) = 84. One-way ANOVA with Dunnett’s test for multiple comparisons in (A-F) and Kruskal-Wallis oneway ANOVA with Dunnett’s test for multiple comparisons in (H). For two group comparisons, twotailed, unpaired Student’s t-test was used in (A-F) and one-tailed, paired Student’s t-test in (H). Box and whisker plots represent maximum value, upper quartile, median, lower quartile and minimum value.

### Colocalization, *in vitro* and direct *in situ* interactions between UNC-104, UNC-17 and MEC-17

To investigate potential interactions between UNC-104, UNC-17, and MEC-17, we examined their expression patterns (under their own promoters) and colocalization within various *C. elegans* neurons. From confocal images it is evident that UNC-104 and UNC-17 colocalize in cholinergic head neurons (map of ACh head neurons: (Pereira et al., 2015)) as well as in cholinergic sublateral neurons (Fig. 5A+B). As expected, UNC-17 expression was not detected in the glutamatergic ALM neuron (e.g., Fig. 5B). UNC-104 and MEC-17 colocalize in the distal part and the branch of the ALM neuron (Fig. 5C) as well as in the soma and proximal part of the ALM neuron (Fig. 5D, single headed arrow). Interestingly, UNC-104/UNC-17 colocalizations improve in *mec-17* knockdown backgrounds (Fig. 5E+G) and UNC-104/MEC-17 colocalizations are impaired in *unc-17* knockdown (Fig. 5F+G) pointing to functional interactions between UNC-104, UNC-17 and MEC-17. This is further validated *in vitro* using co-immunoprecipitation (Co-IP), where interaction between UNC-104 and UNC-17 strengthen when *mec-17* is knocked down and UNC-104/MEC-17 interaction is weakened in case of *unc-17* knockdown (Fig. 5H+I). Future biochemical studies need to reveal the detailed underlying molecular mechanisms of these interactions. Nevertheless, to test for direct protein-protein interactions *in situ*, we employed bimolecular fluorescence assays (BiFC). In BifC assays, the YFP protein Venus is split into two halves (N-terminal Venus, VN, and C-terminal Venus, VC) and the two test-proteins are fused to each half (Fig. 6I). Complementation of Venus, leading to yellow fluorescent signals, occurs only if these two test-proteins are at least 7-10 nm close to each other, therefore likely physically interacting. Critically, it has been shown that interactions are reversible in BiFC, thus, transient interactions are possible to be detected as well (Hsu et al., 2011; Hu & Kerppola, 2003; Kerppola, 2006). From Figure 6 it is obvious that distinct Venus signals appear for the BiFC pairs UNC-104/UNC-17 and UNC-104/MEC-17. Specifically, we show that (similar to our colocalization study) UNC-104/UNC-17 interactions are facilitated in *mec-17* knockdown animals (Fig. 6A-D). Further we reveal successful BiFC signals of UNC-104/MEC-17 in the ALM neuron and mechanosensory neurons in head region (Fig. 6E-H). Moreover, interaction between UNC-104 and MEC-17 was weakened in *mec-17* knockdown animals (Fig. 6D+G-H).

**Figure 5:**
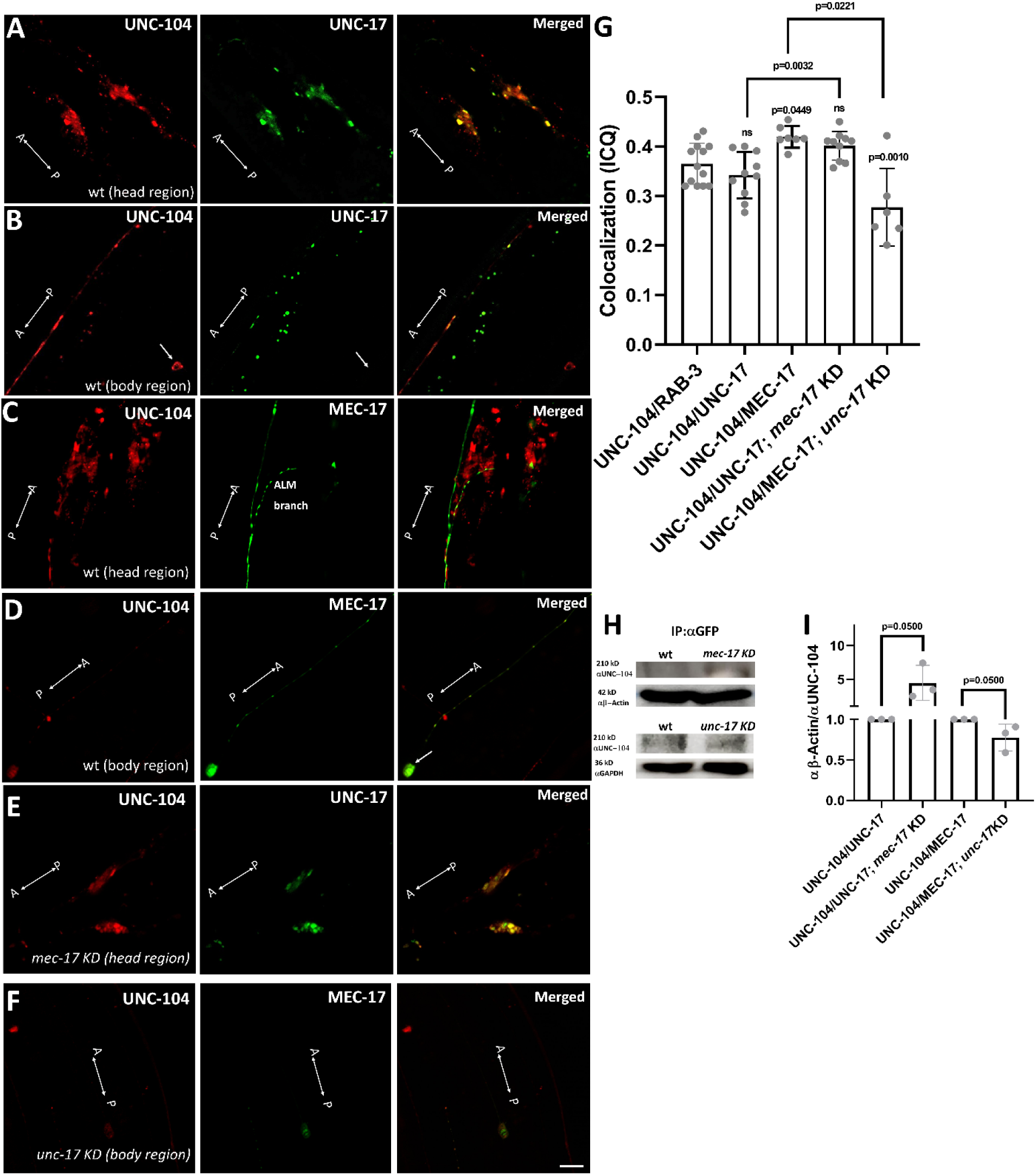
*C. elegans* animals double expressing UNC-104::mRFP/UNC-17::GFP and UNC-104::mRFP/MEC-17::GFP. (A+B) Colocalization between *Punc-104::unc-104::mrfp* and *Punc-17::unc-17::gfp* in (A) the nerve ring and (B) neurons of the (cholinergic) sublateral system (arrow points to an ALM cell body revealing expression of UNC-104 but not UNC-17). (C+D) Colocalization between *Punc-104::unc-104::mrfp* and *Pmec-17::mec-17::gfp* in (C) head neurons (distal ALM and its branch) and (D) body neurons (arrow points to ALM). (E) UNC-104/UNC-17 colocalizations in *mec-17* KD animals. (F) UNC-104/MEC-17 colocalizations in *unc-17* KD animals. (G) Quantification of colocalization using the intensity correlation quotient (ICQ) method. (H+I) Co-IP using anti-GFP antibody to precipitate UNC-104, anti-beta actin antibody to detect β-Actin marker, and anti-GAPDH to detect GAPDH marker. A<->P = anterior/posterior axis. Analyzed worms: 10 per strain. For multiple comparisons, one-way ANOVA with Dunnett’s test. Two-tailed, unpaired Student’s t-test for two group comparisons. Graphs represent mean ± STD. Scale bar: 10 µm.

**Figure 6:**
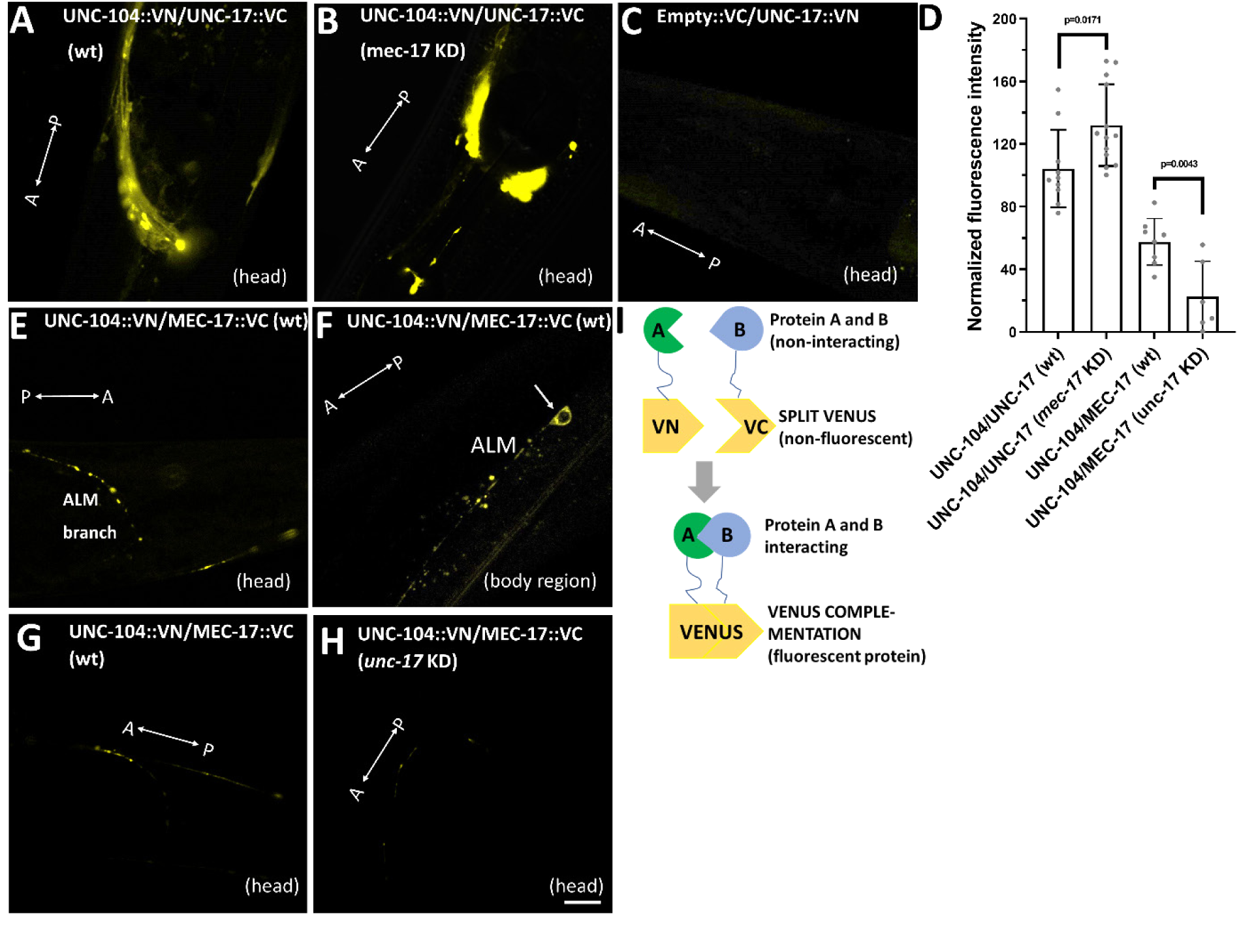
Bimolecular fluorescence complementation (BiFC) assay reveals physical *in situ* interactions between UNC-104/UNC-17 and UNC-104/MEC-17. (A+B) Representative confocal fluorescence images of worms expressing UNC-104::VN/UNC-17::VC in the head of (A) N2 (wt) and (B) in *mec-17* knockdown animals. (C) Empty::VC/UNC-17::VN serving as negative control. (D) Quantification of data shown in (A-B, G-H). (E+F) Worm expressing UNC-104::VN/MEC-17::VC in (E) the head region (showing distal ALM and its branch) and (F) in the body region (showing ALM soma). (G+H) Worms expressing UNC-104::VN/UNC-17::VC in the head of N2(wt) and *unc-17* knockdown animals respectively. (I) Graphical depiction of BiFC assay. A<->P = anterior/posterior axis. Single-headed arrow pointing to ALM soma. Analyzed worms: 10 per strain. Two-tailed, unpaired Student’s t-test. Graphs represent mean ± STD. Scale bar: 10 µm. Scale bar: 10 µm.

## DISCUSSION

### The relation between synaptic transmission defects and UNC-104-based transport

Axonal transport defects have long been linked to various neurodegenerative diseases including HD, AD and PD (Guillaud et al., 2020; Guo et al., 2020; Surana et al., 2020), and the multiple neurological diseases associated to genetic defects in KIF1A/UNC-104 are classified as KAND (KIF1A-associated neurological disorders) encompassing CMT, ALS and HSP (Ali & Yang, 2020; Brady & Morfini, 2017; Millecamps & Julien, 2013). Vice versa, dysfunction or alteration of neurotransmitter release is linked to neuropathological diseases such as AD and PD (Factor et al., 2017; Francis, 2005; Garcia-Alloza et al., 2005). On the other hand, various studies have shown that synaptic transmission defects alter tubulin PTM (Maas et al., 2009) and, conversely, that tubulin PTM regulates axonal transport (Dunn et al., 2008; Peris et al., 2009; Reed et al., 2006; Wloga et al., 2017). The direct link of UNC-104 to synaptic transmission can be seen in a single point mutation (allele *e1265*) in the motor’s C-terminal PH domain (linking the motor to the cargo), causing significant vesicle retention phenotypes that result in a paralyzed and highly uncoordinated nematode (Hall & Hedgecock, 1991; Yonekawa et al., 1998). A homozygous *kif1a* knockout in mouse affects the function of sensory neurons so severely that the animal dies at the embryonic stage (Yonekawa et al., 1998). Similarly, the single amino acid substitution encoded in the hypomorphic allele *unc-17*(*e245*) results in acute neuromuscular function failure leading to paralyzed and coiled-up animals at all developmental stages. In addition, the nematode’s body size is small, pharyngeal pumping is abnormal and locomotion is uncoordinated (de Castro et al., 2009; Sandoval et al., 2006; Zhu et al., 2001). Interestingly, the human ortholog of the unc-17 gene SLC18A3 (solute carrier family 18 member A3) is implicated in AD, HD, pulmonary inflammation and congenital myasthenic syndrome 21 (Banzato et al., 2021; Ferreira-Vieira et al., 2016; Lamond et al., 2021; Smith et al., 2006; Suzuki et al., 2001; Zadrozny et al., 2024). Besides RAB-3 (Fig. 2+3), SNB-1 (synaptobrevin-1) is a well-described cargo for UNC-104 (Bhan et al., 2020; Tien et al., 2011; Wu et al., 2016) and a genetic interaction between UNC-17 and SNB-1 has been reported by others, suggesting that UNC-17 functions together with SNB-1 to maintain proper packaging and loading of ACh (Sandoval et al., 2006). In *unc-17(e245)* mutants, we observed reduced fluorescence intensity of UNC-104 and RAB-3 clusters in ALM neurons, with integrated density decreased by approximately 33.3% for UNC-104 and 37.5% for RAB-3, and mean gray value reduced by approximately 40% for both (Fig. 3I, J). These findings, which complement the reduced cluster size (Fig. 3G, H), indicate impaired aggregation or transport of these proteins, likely due to compromised UNC-104 motility (Fig. 2) driven by increased tubulin acetylation. Additionally, the 35% reduction in UNC-104 mean gray value in *unc-17(e245)* mutants (Fig. 3I) further supports the role of motor dysfunction in these defects. We have shown both colocalization and positive BiFC signals of UNC-104 and UNC-17 in cholinergic head and sublateral neurons (Fig. 5+6) suggesting that UNC-104 associates with synaptic vesicles that transport UNC-17. BiFC assays demonstrate that the interaction between UNC-104 and wild-type UNC-17 is enhanced in a *mec-17* knockdown background, as evidenced by stronger fluorescence signals in cholinergic head neurons (Fig. 6A-D). This indicates that reduced tubulin acetylation, resulting from *mec-17* knockdown, strengthens the UNC-104/UNC-17 interaction, likely by stabilizing the motor-cargo complex on microtubules. Co-IP experiments further confirm this interaction, demonstrating that wild-type UNC-17 robustly co-immunoprecipitates with UNC-104 in a *mec-17* knockdown background, indicating that reduced tubulin acetylation enhances the stability of the UNC-104/UNC-17 complex (Fig. 5H). The effects of ACh agonists such as levamisole (Fig. 4+Suppl. Fig. S5) have been reported by others as well revealing a tight relation between levamisole and RAB-3 motility (Mondal et al., 2011). To avoid that the synaptic transmission defect in unc-17 mutants would negatively affect our observations on how elevated acetylation affects synaptic vesicle transport, we have chosen the glutamatergic ALM neuron for our observations. Because UNC-17 does not express in the ALM neuron, we conclude that the effect of increased tubulin acetylation is non-cell autonomous. Nevertheless, it is interesting that MEC-17 associate with UNC-104 in BiFC assays (Fig. 6) proposing the idea of MEC-17 being a cargo of UNC-104. This interplay between UNC-104, UNC-17, and MEC-17, driven by tubulin acetylation dynamics, is summarized in a model (Fig. 7), which illustrates how increased MT acetylation in *unc-17(e245)* mutants impairs UNC-104 motility and synaptic transmission, with implications for KAND.

**Figure 7:**
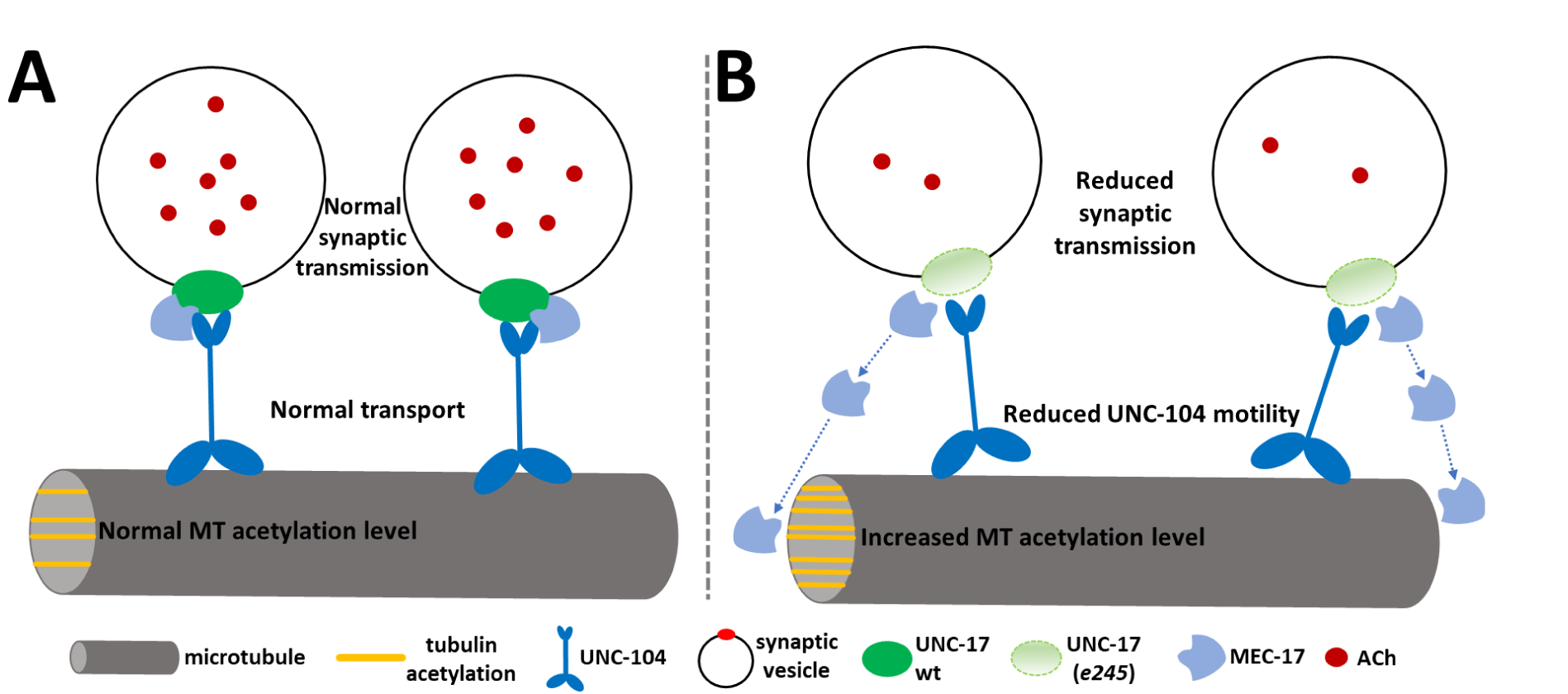
Functional interaction of UNC-104/UNC-17/MEC-17 in neurons. (A) Under normal tubulin acetylation conditions, UNC-104 interacts with UNC-17 and MEC-17 forming a complex while processing along microtubule in cholinergic neurons. (B) When UNC-17 is downregulated, MEC-17 is released from the complex leading to increased tubulin acetylation level affecting the motility of UNC-104 and synaptic transmission.

### The effect of tubulin acetylation on kinesin-1 and kinesin-3 motility in neurons

The consequences of MT acetylation on kinesin-1 motility have been described in various studies. *In vitro* motility assays demonstrate that loss of acetylation on α-tubulin K-40 results in reduced kinesin-1 velocity. In the same study, inhibiting HDAC6 in primary hippocampal cells leads to increased tubulin acetylation levels and concomitantly steering the kinesin-1 cargo JIP1 to the neurite tips (Reed et al., 2006). In cultured neurons and *Drosophila* models with pathogenic mutations of LRRK2 Roc-COR, mitochondrial transport was inhibited in both directions with alongside decreased tubulin acetylation. Increasing tubulin acetylation levels through RNAi knockdown of *HDAC6* and *Sirt2* in mutant *Drosophila* restored the axonal transport of mitochondria and locomotion ability (V. K. Godena et al., 2014). Besides these studies revealing a positive relation between kinesin-1 motility and tubulin acetylation, one study demonstrated reduced (dynein-mediated) aggresome transport in A549 cell lines after *HDAC6* knockdown (leading to increased acetylation levels). Interestingly, interactions between HDAC6 and dynein have been validated (Kawaguchi et al., 2003). Thus, it is not unlikely that PTM enzymes and motors directly interact as revealed for UNC-104 and MEC-17 in the present study (Figure 6E-G). BiFC assays further reveal that the interaction between UNC-104 and wild-type MEC-17 is diminished in an *unc-17* knockdown background, as evidenced by reduced fluorescence signals in head and ALM neurons (Fig. 6D+G-H). This indicates that UNC-17 is critical for maintaining the UNC-104/MEC-17 interaction, potentially by stabilizing the motor-cargo complex or modulating the local microtubule environment through its role in synaptic vesicle transport. Co-IP experiments further reveal that MEC-17 interacts with UNC-104, and in an *unc-17* knockdown background, their interaction is weakened (Fig. 5I). This indicates that UNC-17 plays a critical role in sta-bilizing the UNC-104/MEC-17 interaction, where the presence or absence of UNC-17 influences the interaction between the motor and the acetyltransferase. Due to the ongoing discussion on whether α-tubulin K40 acetylation directly enhances the velocity of kinesin-1 and its affinity to MTs, one group compared kinesin-1 motor motility on MTs reconstituted from acetylated and deacetylated tubulin. As a result, this group has clearly demonstrated that tubulin acetylation alone does not affect kinesin-1 velocity and run length; that it is rather a combination of various PTMs as well as MAP binding (Walter et al., 2012).

In one study, inhibition of HDAC6 (leading to increased acetylation) in primary human macrophages reduced the anterograde velocity and run length of kinesin-3 KIF1C (Bhuwania et al., 2014) underscoring the tendency of motors from the kinesin-3 class to be negatively affected by acetylation (Fig. 4). Furthermore, our findings of functional interactions between a molecular motor (UNC-104), regulator of neurotransmission (UNC-17), and a PTM enzyme (MEC-17) is supported by reports from other groups. For instance, inhibiting glycine receptor activity in primary hippocampal neurons disturbed the transport of gephyrin (that anchors glycine receptors) to the periphery of neurites. Critically, inhibition of glycine receptors significantly increased tubulin polyglutamylation levels, and depletion of polyglutamylase activity with antibodies prevented the inhibition of gephyrin transport (Maas et al., 2009). This study effectively reveals the relation between synaptic transmission, tubulin PTM and axonal transport. In greater relation to our study, both overexpression of MEC-17 and mutations in the *mec-17* gene led to a significant reduction of synaptic vesicles at synapses in *C. elegans* TRNs (Teoh et al., 2022). The Co-IP results (Fig. 5H+I) reinforce this interplay, demonstrating that the UNC-104/UNC-17/MEC-17 complex is dynamically regulated by the presence or absence of each component, with tubulin acetylation playing a pivotal role in stabilizing or destabilizing these interactions In another study, higher MT acetylation levels correlated with the sensitivity to aldicarb (a cholinesterase inhibitor) concomitantly increasing velocities of dense core vesicles in *C. elegans* neurons (Solinger et al., 2010). Moreover, it has been demonstrated that MEC-17 co-distributes with synaptic vesicle precursors in super-resolution images and that MEC-17 localizes to the external surface of vesicles (Even et al., 2019). Critically, it was verified that KIF1A reveals only low affinity to MTs near presynaptic sites. The reason is the enrichment of MT plus-end TIPs (e.g., EB proteins) that increase detachment rates of KIF1A to facilitate proper delivery of precursors to synaptic sites (Guedes-Dias et al., 2019).

### Future outlook

As mentioned above, results from our study point to a non-cell autonomous effect of the *unc-17* mutation to increase tubulin acetylation in the glutamatergic ALM. We therefore assume that modulation of ALM by affected cholinergic upstream neurons must trigger signaling cascades to modulate tubulin PTM. In *C. elegans,* unc-17/VAChT expression defines cholinergic identity and is present in 52 of the 118 classes of adult neurons, amounting to 159 out of 302 neurons (Pereira et al., 2015). Examples of direct input from cholinergic neurons are the head interneurons AVD, AVE, PVC, RIF, RMD, SIA and SDQ (wormatlas.org). Also, more complex circuits need to be considered modulating ALM such as (and among dozens of others): VC -> PVQ -> RMG -> ALM, VC -> VD |-| RIF -> ALM, HSN -> RIF -> ALM or HSN -> BDU -> ALM (with VC and HSN both expressing UNC-17) (wormweb.org). Future studies using laser ablation techniques need to evaluate whether all or only some of these upstream neurons affect tubulin PTM in ALM neurons (or other TRNs).

## ACKNOWLEDGEMENTS

We thank the *C. elegans* Core Facility (CECF) Taiwan (funded by the Ministry of Science and Technology, MOST) for providing microinjection setups and worm observation systems. We acknowledge MOST grant MOST 107-2311-B-007-003- to OIW.

## AUTHOR CONTRIBUTIONS

**CSK** - Data curation, Formal analysis, Writing - review & editing; **MCW** - Data curation, Formal analysis, Writing - review & editing; **OB** - Data curation, Formal analysis, **SNB** - Data curation, Formal analysis; **PB** - Data curation, Formal analysis; **OIW** – Conceptualization, Funding acquisition, Supervision, Writing - review & editing.

## Availability of data

The data that support the findings of this study are available from the corresponding author upon reasonable request.

## MATERIALS & METHODS

### *C. elegans* maintenance, plasmids constructs and generation of strains

Nematodes were maintained at 20°C on NGM agar plates seeded with OP50 *E. coli* according to standard methods (Brenner, 1974). Note that for one experiment (Fig. 4, wt;Levamisole 1mM) we added 10 mM levamisole (#Sigma T1512) to the petri dishes before pouring the (hand warm) NGM agar into them (reaching a final concentration of 1 mM levamisole). If not stated otherwise, strains were obtained from the *Caenorhabditis* Genetic Center (Minnesota, USA). We specifically use monomeric fluorophores for motility studies known to have less aggregation propensities as non-monomeric fluorophores (Campbell et al., 2002). *Punc-104::mcherry::rab-3* was designed by amplifying the rab-3 gene from a cDNA library using primers ATGGCGGCTGGCGGACAA (forward) and TTAGCAATTGCATTGCTGTT (reverse). The resulting plasmid was microinjected (using a standard protocol (Mello et al., 1991)) at 50 ng/μl into either N2 or CB933 *unc-17*(*e245*) strains to generate OIW 118 *N2*;*nthEx118*[*Punc-104*::*mCherry*::*rab-3*] and OIW 119 *unc-17*(*e245*);*nthEx119*[*Punc-104*::*mCherry*::*rab-3*], respectively. The existing plasmid *Punc-104::unc-104::mrfp* (Tien et al., 2011) was injected into *unc-17*(*e245*) strain at 80 ng/μl to generate OIW 120 *unc-17*(*e245*);*nthEx120*[*Punc-104*::*unc-104::mrfp*]. OIW 121 *unc-17(e245);nthEx121[Punc-104::unc-104::mrfp]* was generated by crossing *unc-17(e245)* with existing strain *unc-104(e1265);nthEx[Punc-104::unc-104::mrfp]* (Tien et al., 2011) and the strong coiler phenotype of *unc-17*(*e245*) was used for screening after backcrossing. Note that for optimal operation of the Punc104 promoter (thus optimal UNC-104 expression) rather high injection dosages are needed (Tien et al., 2011; Wagner et al., 2009). Indeed, only unc-104 (full-length) plasmid concentrations above 70 ng/µl fully rescue the highly uncoordinated *unc-104*(*e1265*) phenotype (Klopfenstein & Vale, 2004) and neither ectopic expression nor any effects on worm behavior are detected. To generate the transcriptional fusion plasmid *Punc-17::gfp*, the 1.9 kb promoter region of *unc-17* was cloned into the existing pPD95.77 GFP vector plasmid taken from the Andrew Fire Library (a kind gift from Dr. Wu Yi-Chun at NTU) using the aaaaaGCATGCAATGACTAGACGCTGAAG (forward) and aaaaaCTGCAGGAAAATTAAATATTTTAGTGTAAAACTTTTGGG (reverse) primers. To generate the translational fusion plasmid *Punc-17::unc-17::gfp*, the cDNA of *unc-17* was cloned into the transcriptional fusion plasmid *Punc-17::gfp* using aaaaaaTCTAGAATGGGCTTCAACGTGCCCGTC (forward) and aaaaaaGGTACCCTACCACTGCGGATTCAGTGG (reverse) primers. OIW122 *N2;nthEx122[Punc-17::unc-17::gfp;unc-104::unc-104::mrfp]* was generated by microinjecting the *Punc-17::unc-17::gfp* plasmid (80 ng/μl) and *Punc-104::unc-104::mrfp* plasmid (80 ng/μl) into wild type N2 worms for colocalization analysis. The translational fusion plasmid *Pmec-17::mec-17::gfp* was generated by cloning the promoter region of *mec-17* into the existing pPD95.77 GFP vector plasmid using primer aaaaGCATGCgataagaattggtagagcaaagacggc (forward) and aaaaGTCGACgatcgaatcgtctcacaactgatcca (reverse) primers. The genomic DNA of *mec-17* was then cloned into the plasmid using aaaaaCCCGGGatgcaagtcgacgccgacc (forward) and aaaaaTGGCCAaccacaatggcttgctcgac (reverse) primers. To observe MEC-17 overexpression effects, OIW 123 *N2;nthEx123[Pmec-17::mec-17::gfp;Punc-104::unc-104::mrfp]* was generated by microinjecting the *Pmec-17::mec-17::gfp* plasmid (100 ng/μl) and *Punc-104::unc-104::mrfp* plasmid (90 ng/μl) into N2 wild type worms. For BiFC assays, *Punc-17::unc-17::VC155* and *Punc-17::unc-17::VN173* were generated by subcloning the promoter and cDNA of *unc-17* from aforementioned *Punc-17::unc-17::gfp* into the existing plasmids *Punc-104::VC155* and *Punc-104::VN173* (Hsu et al., 2011), respectively. *Pmec-17::mec-17::VC155* was generated by subcloning the promoter and genomic DNA of *mec-17* from *Pmec-17::mec-17::gfp* into the existing plasmid *Punc-104::VC155*. OIW 124 *N2;nthEx124[Punc-104::unc-104::VN173;Punc-17::unc-17::VC155]* was then generated by microinjecting existing plasmid *Punc-104::unc-104::VN173* (60 ng/μl) (Wu et al., 2016) and above mentioned *Punc-17::unc-17::VC155* (60 ng/μl) plasmids along with roller co-injection marker pRF4(rol-6(su1006)) (50 ng/μl) into N2 worms. OIW 125 *N2;nthEx125[Punc-104::unc-104::VN173;Pmec-17::mec-17::VC155]* was generated by microinjecting *Punc-104::unc-104::VN173* (60 ng/μl) and plasmid *Pmec-17::mec-17::VC155* (60 ng/μl) plasmids along with roller co-injection marker pRF4(rol-6(su1006)) (50 ng/μl) into N2 worms. For a negative control, we injected the *Punc-104::VC155* empty vector plasmid (40 ng/μl) and *Punc-17::unc-17::VN173* plasmid (60 ng/μl) along with roller co-injection marker pRF4(rol-6(su1006)) (50 ng/μl) into N2 worms to generate OIW 126 *N2;nthEx127[Punc-104::VC155EV; Punc-17::unc-17::VN173]*.

### RNAi feeding assay

For RNA interference experiments, we employed the RNAi feeding method (Kamath et al., 2001) in which worms are fed by bacteria producing the desired dsRNA. Feeding clones were obtained from Julie Ahringer’s *C. elegans* RNAi feeding library (Source BioScience LifeSciences, USA; a kind gift from the *C. elegans* Core Facility, Taiwan) and sequenced them to determine their correctness. NGM plates containing ampicillin (100 μg/ml) and 1 mM IPTG were inoculated with the respective HT115 *E. coli* strain carrying the appropriate gene insert (flanked by T7 promoters) and grown overnight. 15-20 worms were transferred to the respective RNAi feeding plate and incubated at 20°C. Worms were then transferred to new RNAi feeding plates every 24 hours and the F1 progeny was used for analysis after day 5. Note that knocking down proteins in neuronal *C. elegans* tissues without the use of “enhancer strains” (such as *rrf-3* or *eri-1*) has been successfully implemented in various studies from this lab (Barmaver et al., 2022; Bhan et al., 2020; Chen et al., 2019; Muniesh et al., 2020; Wu et al., 2016).

### Western blot and co-immunoprecipitation assays

To perform Western blotting from whole worm lysates, we employed a published protocol (Mahmood & Yang, 2012). Here, 100 µg sample protein is used and membrane blocking is done with milk powder for 2 hours at room temperature (RT). We use primary anti-acetylated tubulin antibody from mouse (monoclonal 6-11 B-1, T7451 Sigma, 1/1000 dilution) as well as primary anti-tyrosinated tubulin antibody from mouse (monoclonal TUB-1A2, T9028 Sigma, 1/5000 dilution) along with mouse anti-GAPDH antibody (monoclonal 60004-1-Ig, Proteintech, 1/5000 dilution) at 4°C overnight. The secondary anti-mouse antibody (GTX213111-01, GeneTex) was used at 1/2000 dilution for acetylation, 1/5000 dilution for tyrosination, and 1/5000 dilution for GAPDH detection and incubated at RT for 2 hours. For Co-IP, 2μg of anti-GFP antibody (GTX628528, GeneTex) was added to 1 mg of protein samples to pull down UNC-104 using protein G magnetic beads (LSKMAGG10, Millipore) and incubated with IP buffer at 4°C for overnight. Target proteins were then detected by Western blotting using primary antibodies anti-UNC-104 (commissioned by our group, polyclonal, rabbit, GeneTex, 1/500 dilution) to detect MEC-17 or UNC-17 interacting with UNC-104, anti-beta Actin (polyclonal, rabbit, GTX109639, GeneTex, 1/2500 dilution), and anti-GAPDH (polyclonal, rabbit, GTX100118, GeneTex, 1/5000 dilution). Secondary antibody anti-rabbit (polyclonal, GTX213110-01, GeneTex) was used at 1/1000 dilution for UNC-104, and 1/5000 dilution for beta Actin and GAPDH detection. ECL (CyECL Western Blotting Substrate H) was applied for chemiluminescent detection and the membrane was incubated in ECL solution. We used a LAS-4000 Luminescent image analyzer to develop the blots. All experiments were performed in triplicate and quantified after normalization using NIH ImageJ software.

### Whole mount immunostaining

For whole-mount immunostaining, we followed a published protocol (Duerr, 2006). Nematodes are treated with 4% paraformaldehyde in a 2 ml tube and then incubated at 37°C in beta-mercaptoethanol solution for 2 hours. Collagenase is then added to the solution and worms are incubated for an additional 6.5 hours at 37°C. For antibody staining, worms are blocked first for 1 hour at RT using blocking buffer (5% lyophilized serum albumin solution and antibody buffer) and after applying primary antibodies (anti-acetylated 6-11 B-1 1:200) incubated at 4°C overnight. Secondary antibodies (antimouse IgG CF488, Sigma) are then used for 2-4 hours at RT and the solution is covered by aluminum foil to protect the worms from light avoiding photobleaching. 10 µl of the sample is then mixed with 10 µl of mounting medium (200 mg n-propyl gallate, 0.3 ml 1 M Tris pH9, 7 ml glycerol, and 2.7 ml ddH_2_O) on poly-L-lysine coated glass slide (Sigma) covered with a 22 mm x 22 mm cover slide. A Zeiss LSM780 confocal microscope was employed for imaging worms and ImageJ to determine “corrected total cell fluorescence” with CTCF = Integrated density of selection region – (Area of selection region x Mean fluorescence of background).

### Worm imaging and motility analysis

For microscopic observations, worms were immobilized on 2% agarose-coated cover slides, and no anesthetics (such as levamisole) were used. A Zeiss LSM780 confocal laser scanning microscope was employed for imaging worms as shown in Figures 5 and 6. For motility analysis (Figs. 2 + 4), we employed an Olympus IX81 microscope with a DSU Nipkow spinning disk unit connected to an Andor iXon DV897 EMCCD camera for high-speed and long-duration time-lapse imaging (4-5 frames per second). To convert recorded time-lapse sequences into kymographs, we use the imaging analysis software NIH ImageJ. The ‘straighten plugin’ is employed to straighten curved axons, and after drawing a line over the axon, the plugin ‘reslice stack function’ is executed to generate kymographs. In kymographs, static particles appear as vertical lines, whereas the slope of a moving particle corresponds to the velocity (speed) of the particle (Fig. 2E). A pause is defined if motors move less than 0.07 μm/s, and each calculated velocity event does not contain any pauses. A moving event is defined as a single motility occurrence typically right after a pause or a reversal, and such an event ends when the motor again pauses or reverses. Anterograde and retrograde run lengths represent the distances traveled by particles in their respective directions within the kymograph. Anterograde TTRL and retrograde TTRL were calculated by summing all forward or backward displacements, respectively, during the observation period. Distance traveled was obtained by adding both anterograde and retrograde run lengths, reflecting the cumulative movement of the particle regardless of direction. Note that data shown in Fig. 2 and Fig. 4 were acquired independently by two students (with experiments spaced apart by five years), thus the effect of *unc-17*(*e245*) on UNC-104 motility was well reproduced. Motor cluster analysis (Fig. 3) was carried out using ImageJ’s ‘area’ tool and the ‘particle analyze’ plugin. To measure travel distances (Fig. 3) of particles in (straightened) neurons, we used ImageJ’s ‘line tool’ to draw a line from the proximal axon hillock to the identified distal particle (Fig. 5A). Intensity correlation quotient (ICQ) (Fig. 5E) was determined by selecting the region of interest using the ‘polygonal selection tool’ in ImageJ. After background fluorescent intensity subtraction (‘subtract background’ function), the plugin ‘intensity correlation analysis’ was used to generate ICQ values. ICQ values range from -0.5 to 0.5 with values close to 0.5 indicating interdependent expression of two fluorophores, values close to 0 implying random expression and values close to -0.5 revealing segregated expression.

### BiFC assays

Bimolecular fluorescence complementation assays (BiFC) are similar to “split-GFP” experiments. Here, a yellow fluorescent protein (YFP) VENUS is split into two halves: a N-terminal part VN and a C-terminal part VC. Both parts are non-fluorescent but can complement (when close enough to each other) to generate a fully functional VENUS (see Fig. 6H). Each part is then tagged to one of the test-proteins. When the two test-proteins are close enough (about 7-10 nm apart), fluorescence complementation may occur suggesting physical interactions between the two test-proteins (Fan et al., 2008; Hsu et al., 2011; Shyu et al., 2008). Cloning of BiFC constructs used for this study is discussed above.

### Statistics

To understand whether analyzed data distributions follow normal (Gaussian) distributions (one- or two tailed), we blot histograms in the first place. In case of present normal distributions, we use parametric tests and in case of non-normal distributions we apply non-parametric tests. For parametric tests, we use one-way ANOVA with Dunnett’s test for multiple-group comparisons and Student’s t-test for two-group comparisons. For Student’s t-test we use unequal variance and decide one- or two-tailed distributions based on blotted histograms. In this study, we only used a non-parametric test (Kruskal-Wallis one-way ANOVA with Dunnett’s test for multiple comparisons) for Fig. 4H in which seven wild type data points are all the same value (due to the design of this method) not allowing for a parametric test. Note that for Fig. 1B and S1 we still use parametric tests due to small sample size (three data points) allowing us to test for a normal distribution null-hypothesis.

---

**Fig. S1:**
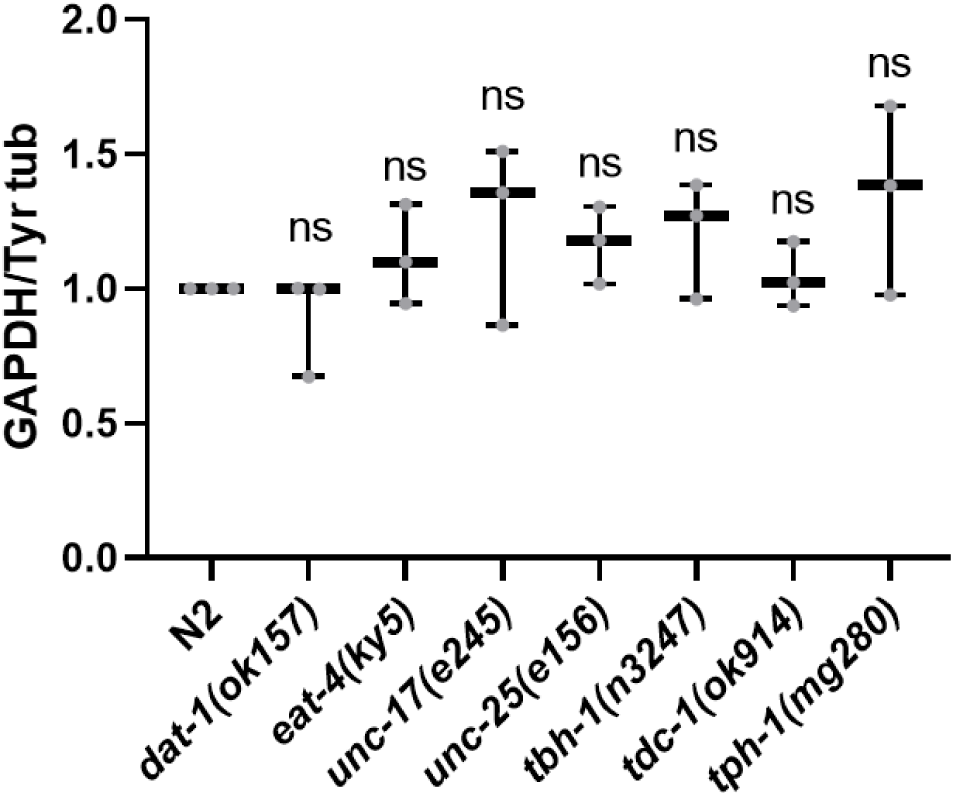
Quantification of band signal intensity from αTyr tubulin gel as shown in Figure 1A, lower panel. One-way ANOVA with Dunnett’s test for multiple comparisons utilized with *p<0.05. Box and whisker plots represent maximum value, upper quartile, median, lower quartile and minimum value. Scale bars: 10 µm.

**Fig. S2:**
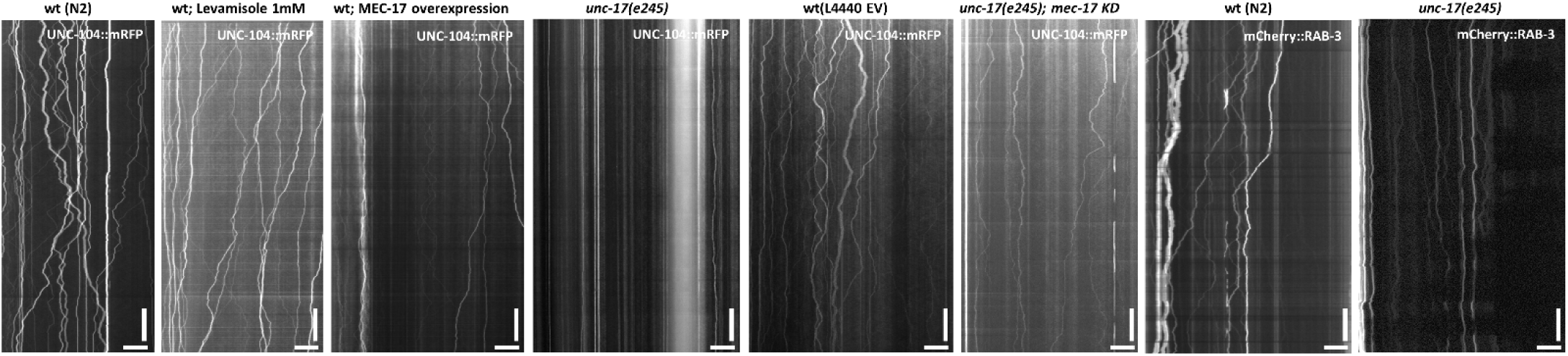
Representative kymographs employed to generate data as shown in Fig. 2+4.

**Fig. S3:**
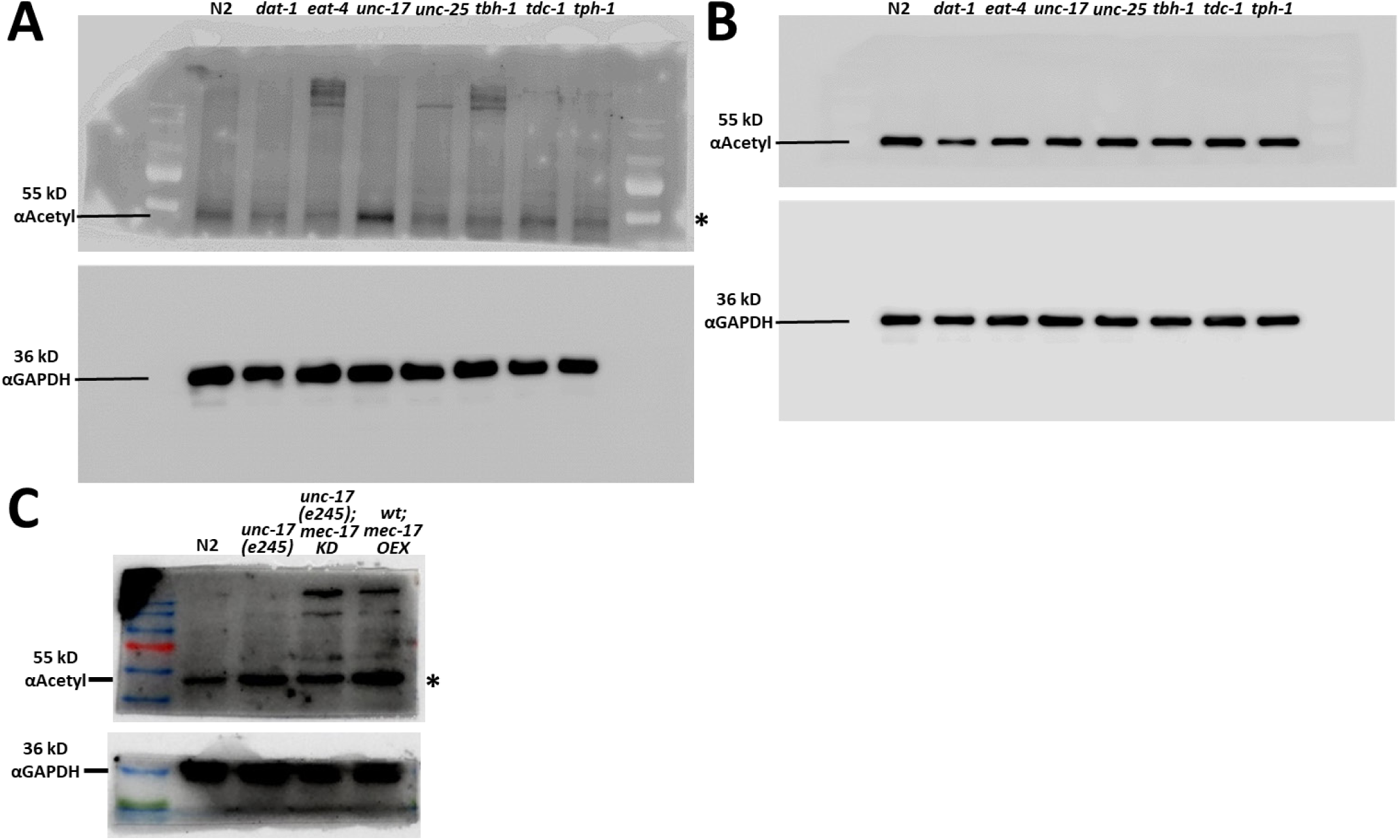
Raw images for cropped gels shown in Figs. 1A and 4G. (A) Upper gel is αAceyl tubulin (55kD) and lower gel is αGAPDH (36kD) as shown in Figure 1A (upper panel). (B) Upper gel is αTyrosin tubulin (55kD) and lower gel is αGAPDH (36kD) as shown in Figure 1A (lower panel). (C) Upper gel is αAceyl tubulin (55kD) and lower gel is αGAPDH (36kD) as shown in Figure 4G. Wildcard marks bands specific to the antibody.

**Fig. S4:**
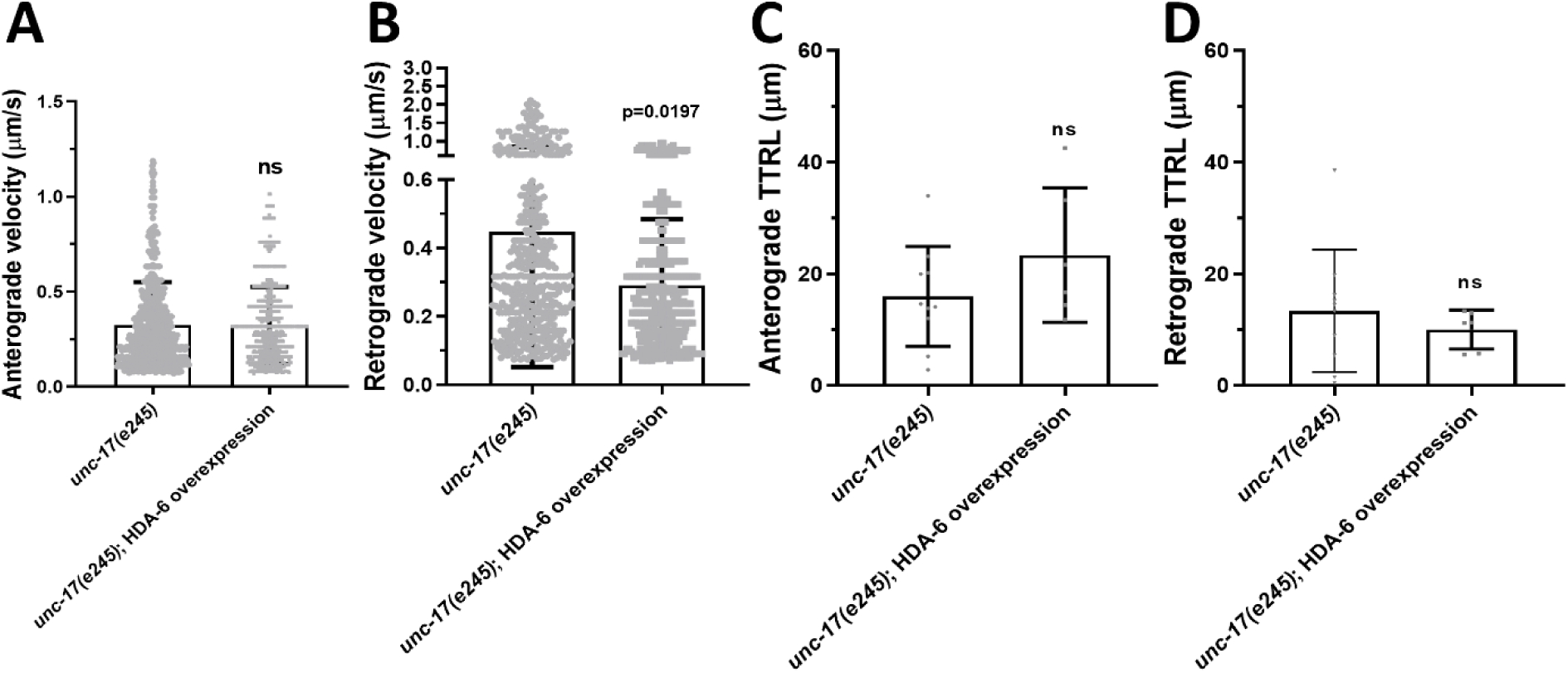
Motor motility after overexpression of deacetylase. Quantification of various motility parameters of UNC-104::mRFP in neurons of living C. elegans animals. Deacetylase HAD-6 was overexpressed (OEX) in *unc-17* (e245) animals. (A) Anterograde velocity. (B) Retrograde velocity. (C) Total anterograde and (D) Retrograde run lengths. Two-tailed, unpaired Student’s t-test

**Fig. S5:**
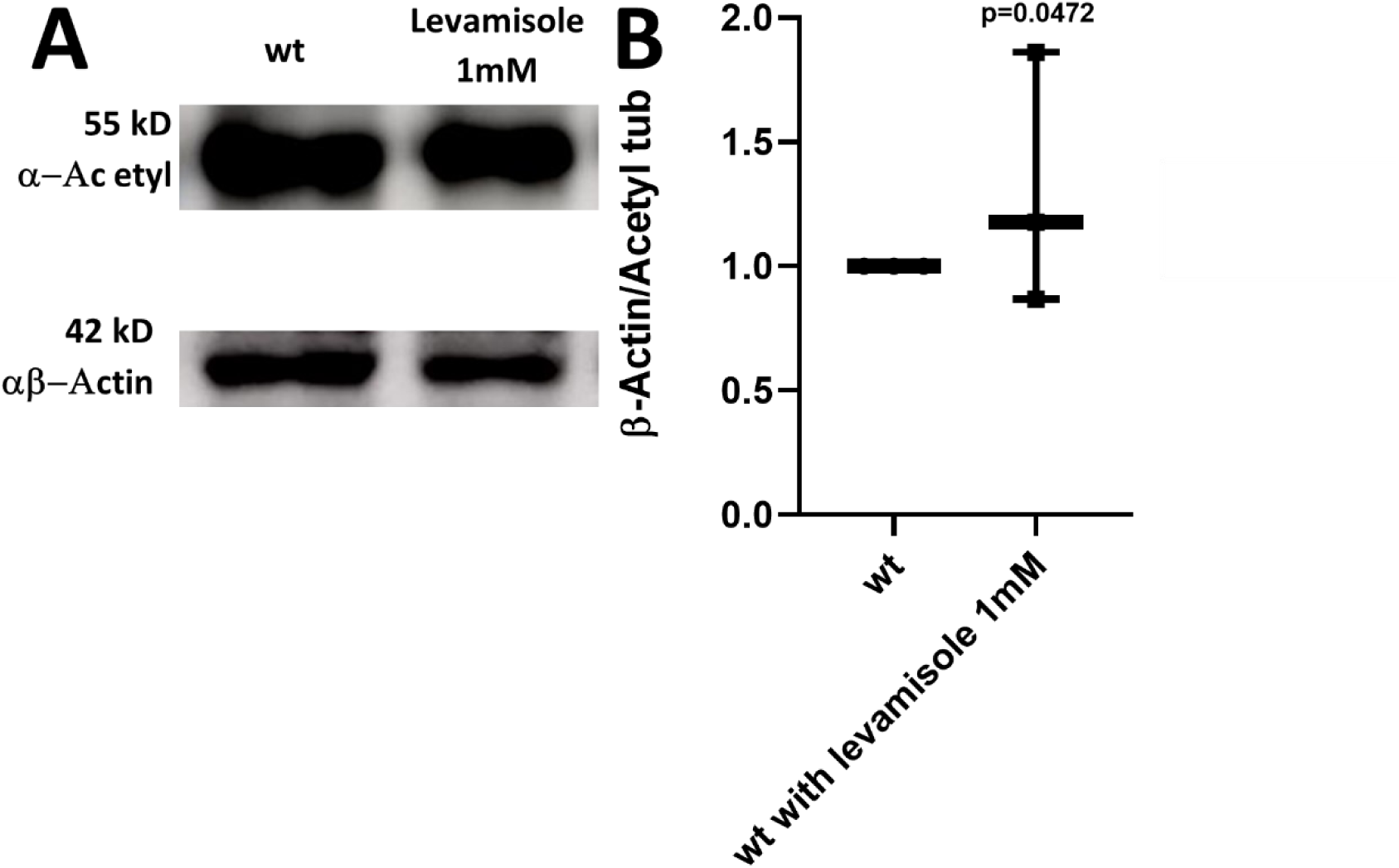
Effect of Ach transmission inhibitor in acetylation level. (A) Western blot revealing levamisole (1mM) effect on tubulin acetylation level. (B) Quantification of band intensity of gel shown in (A). One sample t-test.

## REFERENCES

1. Alfonso, A., Grundahl, K., Duerr, J. S., Han, H. P., & Rand, J. B. (1993). The Caenorhabditis elegans unc-17 gene: a putative vesicular acetylcholine transporter. Science, 261(5121), 617–619. 10.1126/science.8342028

2. Ali, I., & Yang, W. C. (2020). The functions of kinesin and kinesin-related proteins in eukaryotes. Cell Adh Migr, 14(1), 139–152. 10.1080/19336918.2020.1810939

3. Alkema, M. J., Hunter-Ensor, M., Ringstad, N., & Horvitz, H. R. (2005). Tyramine Functions independently of octopamine in the Caenorhabditis elegans nervous system. Neuron, 46(2), 247–260. 10.1016/j.neuron.2005.02.024

4. Avery, L. (1993). The genetics of feeding in Caenorhabditis elegans. Genetics, 133(4), 897–917. https://www.ncbi.nlm.nih.gov/pubmed/8462849

5. Banzato, R., Pinheiro, N. M., Olivo, C. R., Santana, F. R., Lopes, F., Caperuto, L. C., Câmara, N. O., Martins, M. A., Tibério, I., Prado, M. A. M., Prado, V. F., & Prado, C. M. (2021). Long-term endogenous acetylcholine deficiency potentiates pulmonary inflammation in a murine model of elastase-induced emphysema. Sci Rep, 11(1), 15918. 10.1038/s41598-021-95211-3

6. Barmaver, S. N., Muthaiyan Shanmugam, M., Chang, Y., Bayansan, O., Bhan, P., Wu, G. H., & Wagner, O. I. (2022). Loss of intermediate filament IFB-1 reduces mobility, density, and physiological function of mitochondria in Caenorhabditis elegans sensory neurons. Traffic, 23(5), 270–286. 10.1111/tra.12838

7. Bhan, P., Muthaiyan Shanmugam, M., Wang, D., Bayansan, O., Chen, C. W., & Wagner, O. I. (2020). Characterization of TAG-63 and its role on axonal transport in C. elegans. Traffic, 21(2), 231–249. 10.1111/tra.12706

8. Bhuwania, R., Castro-Castro, A., & Linder, S. (2014). Microtubule acetylation regulates dynamics of KIF1C-powered vesicles and contact of microtubule plus ends with podosomes. Eur J Cell Biol, 93(10-12), 424–437. 10.1016/j.ejcb.2014.07.006

9. Binotti, B., Jahn, R., & Chua, J. J. (2016). Functions of Rab Proteins at Presynaptic Sites. Cells, 5(1). 10.3390/cells5010007

10. Brady, S. T., & Morfini, G. A. (2017). Regulation of motor proteins, axonal transport deficits and adult-onset neurodegenerative diseases. Neurobiol Dis, 105, 273–282. 10.1016/j.nbd.2017.04.010

11. Brenner, S. (1974). The genetics of Caenorhabditis elegans. Genetics, 77(1), 71–94. http://www.genetics.org/content/77/1/71.full.pdf

12. Campbell, R. E., Tour, O., Palmer, A. E., Steinbach, P. A., Baird, G. S., Zacharias, D. A., & Tsien, R. Y. (2002). A monomeric red fluorescent protein. Proc Natl Acad Sci U S A, 99(12), 7877–7882. 10.1073/pnas.082243699

13. Chen, C. W., Peng, Y. F., Yen, Y. C., Bhan, P., Muthaiyan Shanmugam, M., Klopfenstein, D. R., & Wagner, O. I. (2019). Insights on UNC-104-dynein/dynactin interactions and their implications on axonal transport in Caenorhabditis elegans. J Neurosci Res, 97(2), 185–201. 10.1002/jnr.24339

14. Chiba, K., Kita, T., Anazawa, Y., & Niwa, S. (2023). Insight into the regulation of axonal transport from the study of KIF1A-associated neurological disorder. J Cell Sci, 136(5). 10.1242/jcs.260742

15. Cueva, J. G., Hsin, J., Huang, K. C., & Goodman, M. B. (2012). Posttranslational acetylation of alpha-tubulin constrains protofilament number in native microtubules. Curr Biol, 22(12), 1066–1074. 10.1016/j.cub.2012.05.012

16. d’Ydewalle, C., Krishnan, J., Chiheb, D. M., Van Damme, P., Irobi, J., Kozikowski, A. P., Vanden Berghe, P., Timmerman, V., Robberecht, W., & Van Den Bosch, L. (2011). HDAC6 inhibitors reverse axonal loss in a mouse model of mutant HSPB1-induced Charcot-Marie-Tooth disease [Research Support, Non-U.S. Gov’t]. Nat Med, 17(8), 968–974. 10.1038/nm.2396

17. de Castro, B. M., De Jaeger, X., Martins-Silva, C., Lima, R. D., Amaral, E., Menezes, C., Lima, P., Neves, C. M., Pires, R. G., Gould, T. W., Welch, I., Kushmerick, C., Guatimosim, C., Izquierdo, I., Cammarota, M., Rylett, R. J., Gomez, M. V., Caron, M. G., Oppenheim, R. W., … Prado, V. F. (2009). The vesicular acetylcholine transporter is required for neuromuscular development and function. Mol Cell Biol, 29(19), 5238–5250. 10.1128/MCB.00245-09

18. Dompierre, J. P., Godin, J. D., Charrin, B. C., Cordelieres, F. P., King, S. J., Humbert, S., & Saudou, F. (2007). Histone deacetylase 6 inhibition compensates for the transport deficit in Huntington’s disease by increasing tubulin acetylation. J Neurosci, 27(13), 3571–3583. 10.1523/JNEUROSCI.0037-07.2007

19. Duerr, J. S. (2006). Immunohistochemistry. WormBook, 1–61. 10.1895/wormbook.1.105.1

20. Dunn, S., Morrison, E. E., Liverpool, T. B., Molina-Paris, C., Cross, R. A., Alonso, M. C., & Peckham, M. (2008). Differential trafficking of Kif5c on tyrosinated and detyrosinated microtubules in live cells. J Cell Sci, 121(Pt 7), 1085–1095. 10.1242/jcs.026492

21. Eshun-Wilson, L., Zhang, R., Portran, D., Nachury, M. V., Toso, D. B., Lohr, T., Vendruscolo, M., Bonomi, M., Fraser, J. S., & Nogales, E. (2019). Effects of alpha-tubulin acetylation on microtubule structure and stability. Proc Natl Acad Sci U S A, 116(21), 10366–10371. 10.1073/pnas.1900441116

22. Even, A., Morelli, G., Broix, L., Scaramuzzino, C., Turchetto, S., Gladwyn-Ng, I., Le Bail, R., Shilian, M., Freeman, S., Magiera, M. M., Jijumon, A. S., Krusy, N., Malgrange, B., Brone, B., Dietrich, P., Dragatsis, I., Janke, C., Saudou, F., Weil, M., & Nguyen, L. (2019). ATAT1-enriched vesicles promote microtubule acetylation via axonal transport. Sci Adv, 5(12), eaax2705. 10.1126/sciadv.aax2705

23. Factor, S. A., McDonald, W. M., & Goldstein, F. C. (2017). The role of neurotransmitters in the development of Parkinson’s disease-related psychosis. Eur J Neurol, 24(10), 1244–1254. 10.1111/ene.13376

24. Fan, J. Y., Cui, Z. Q., Wei, H. P., Zhang, Z. P., Zhou, Y. F., Wang, Y. P., & Zhang, X. E. (2008). Split mCherry as a new red bimolecular fluorescence complementation system for visualizing protein-protein interactions in living cells. Biochem Biophys Res Commun, 367(1), 47–53. S0006-291X(07)02706-4 [pii] 10.1016/j.bbrc.2007.12.101

25. Fan, R., & Lai, K. O. (2022). Understanding how kinesin motor proteins regulate postsynaptic function in neuron. FEBS J, 289(8), 2128–2144. 10.1111/febs.16285

26. Ferreira-Vieira, T. H., Guimaraes, I. M., Silva, F. R., & Ribeiro, F. M. (2016). Alzheimer’s disease: Targeting the Cholinergic System. Curr Neuropharmacol, 14(1), 101–115. 10.2174/1570159x13666150716165726

27. Francis, P. T. (2005). The interplay of neurotransmitters in Alzheimer’s disease. CNS Spectr, 10(11 Suppl 18), 6–9. https://www.ncbi.nlm.nih.gov/pubmed/16273023

28. Garcia-Alloza, M., Gil-Bea, F. J., Diez-Ariza, M., Chen, C. P., Francis, P. T., Lasheras, B., & Ramirez, M. J. (2005). Cholinergic-serotonergic imbalance contributes to cognitive and behavioral symptoms in Alzheimer’s disease. Neuropsychologia, 43(3), 442–449. 10.1016/j.neuropsychologia.2004.06.007

29. Genova, M., Grycova, L., Puttrich, V., Magiera, M. M., Lansky, Z., Janke, C., & Braun, M. (2023). Tubulin polyglutamylation differentially regulates microtubule-interacting proteins. EMBO J, 42(5), e112101. 10.15252/embj.2022112101

30. Godena, V. K., Brookes-Hocking, N., Moller, A., Shaw, G., Oswald, M., Sancho, R. M., Miller, C. C. J., Whitworth, A. J., & De Vos, K. J. (2014). Increasing microtubule acetylation rescues axonal transport and locomotor deficits caused by LRRK2 Roc-COR domain mutations. Nature Communications, 5(1), 5245. 10.1038/ncomms6245

31. Guedes-Dias, P., Nirschl, J. J., Abreu, N., Tokito, M. K., Janke, C., Magiera, M. M., & Holzbaur, E. L. F. (2019). Kinesin-3 Responds to Local Microtubule Dynamics to Target Synaptic Cargo Delivery to the Presynapse. Curr Biol, 29(2), 268–282 e268. 10.1016/j.cub.2018.11.065

32. Guillaud, L., El-Agamy, S. E., Otsuki, M., & Terenzio, M. (2020). Anterograde Axonal Transport in Neuronal Homeostasis and Disease. Front Mol Neurosci, 13, 556175. 10.3389/fnmol.2020.556175

33. Guo, W., Stoklund Dittlau, K., & Van Den Bosch, L. (2020). Axonal transport defects and neurodegeneration: Molecular mechanisms and therapeutic implications. Semin Cell Dev Biol, 99, 133–150. 10.1016/j.semcdb.2019.07.010

34. Hall, D. H., & Hedgecock, E. M. (1991). Kinesin-related gene unc-104 is required for axonal transport of synaptic vesicles in C. elegans. Cell, 65(5), 837–847. https://doi.org/0092-8674(91)90391-B [pii]

35. Hancock, W. O. (2014). Bidirectional cargo transport: moving beyond tug of war. Nat Rev Mol Cell Biol, 15(9), 615–628. 10.1038/nrm3853

36. Hirokawa, N., Niwa, S., & Tanaka, Y. (2010). Molecular motors in neurons: transport mechanisms and roles in brain function, development, and disease. Neuron, 68(4), 610–638. S0896-6273(10)00781-6 [pii] 10.1016/j.neuron.2010.09.039

37. Hoerndli, F. J., Maxfield, D. A., Brockie, P. J., Mellem, J. E., Jensen, E., Wang, R., Madsen, D. M., & Maricq, A. V. (2013). Kinesin-1 regulates synaptic strength by mediating the delivery, removal, and redistribution of AMPA receptors. Neuron, 80(6), 1421–1437. 10.1016/j.neuron.2013.10.050

38. Hsu, C. C., Moncaleano, J. D., & Wagner, O. I. (2011). Sub-cellular distribution of UNC-104(KIF1A) upon binding to adaptors as UNC-16(JIP3), DNC-1(DCTN1/Glued) and SYD-2(Liprin-alpha) in C. elegans neurons. Neuroscience, 176, 39–52. 10.1016/j.neuroscience.2010.12.044

39. Hu, C. D., & Kerppola, T. K. (2003). Simultaneous visualization of multiple protein interactions in living cells using multicolor fluorescence complementation analysis. Nat Biotechnol, 21(5), 539–545. http://www.ncbi.nlm.nih.gov/entrez/query.fcgi?cmd=Retrieve&db=PubMed&dopt=Citation&list_uids=12692560

40. Ikegami, K., Heier, R. L., Taruishi, M., Takagi, H., Mukai, M., Shimma, S., Taira, S., Hatanaka, K., Morone, N., Yao, I., Campbell, P. K., Yuasa, S., Janke, C., Macgregor, G. R., & Setou, M. (2007). Loss of alpha-tubulin polyglutamylation in ROSA22 mice is associated with abnormal targeting of KIF1A and modulated synaptic function. Proc Natl Acad Sci U S A, 104(9), 3213–3218. http://www.ncbi.nlm.nih.gov/entrez/query.fcgi?cmd=Retrieve&db=PubMed&dopt=Citation&list_uids=17360631

41. Janke, C., & Magiera, M. M. (2020). The tubulin code and its role in controlling microtubule properties and functions. Nat Rev Mol Cell Biol, 21(6), 307–326. 10.1038/s41580-020-0214-3

42. Kamath, R. S., Martinez-Campos, M., Zipperlen, P., Fraser, A. G., & Ahringer, J. (2001). Effectiveness of specific RNA-mediated interference through ingested double-stranded RNA in Caenorhabditis elegans. Genome Biol, 2(1), RESEARCH0002. 10.1186/gb-2000-2-1-research0002

43. Kaul, N., Soppina, V., & Verhey, K. J. (2014). Effects of alpha-tubulin K40 acetylation and detyrosination on kinesin-1 motility in a purified system. Biophys J, 106(12), 2636–2643. 10.1016/j.bpj.2014.05.008

44. Kawaguchi, Y., Kovacs, J. J., McLaurin, A., Vance, J. M., Ito, A., & Yao, T. P. (2003). The deacetylase HDAC6 regulates aggresome formation and cell viability in response to misfolded protein stress. Cell, 115(6), 727–738. 10.1016/s0092-8674(03)00939-5

45. Kern, J. V., Zhang, Y. V., Kramer, S., Brenman, J. E., & Rasse, T. M. (2013). The kinesin-3, unc-104 regulates dendrite morphogenesis and synaptic development in Drosophila. Genetics, 195(1), 59–72. 10.1534/genetics.113.151639

46. Kerppola, T. K. (2006). Design and implementation of bimolecular fluorescence complementation (BiFC) assays for the visualization of protein interactions in living cells. Nat Protoc, 1(3), 1278–1286. http://www.ncbi.nlm.nih.gov/entrez/query.fcgi?cmd=Retrieve&db=PubMed&dopt=Citation&list_uids=17406412

47. Klopfenstein, D. R., & Vale, R. D. (2004). The lipid binding pleckstrin homology domain in UNC-104 kinesin is necessary for synaptic vesicle transport in Caenorhabditis elegans. Mol Biol Cell, 15(8), 3729–3739. http://www.ncbi.nlm.nih.gov/entrez/query.fcgi?cmd=Retrieve&db=PubMed&dopt=Citation&list_uids=15155810

48. Lamond, A., Buckley, D., O’Dea, J., & Turner, L. (2021). Variants of SLC18A3 leading to congenital myasthenic syndrome in two children with varying presentations. BMJ Case Rep, 14(1). 10.1136/bcr-2020-237799

49. Li, L., & Yang, X. J. (2015). Tubulin acetylation: responsible enzymes, biological functions and human diseases. Cell Mol Life Sci, 72(22), 4237–4255. 10.1007/s00018-015-2000-5

50. Li, L. B., Lei, H., Arey, R. N., Li, P., Liu, J., Murphy, C. T., Xu, X. Z., & Shen, K. (2016). The Neuronal Kinesin UNC-104/KIF1A Is a Key Regulator of Synaptic Aging and Insulin Signaling-Regulated Memory. Curr Biol, 26(5), 605–615. 10.1016/j.cub.2015.12.068

51. Maas, C., Belgardt, D., Lee, H. K., Heisler, F. F., Lappe-Siefke, C., Magiera, M. M., van Dijk, J., Hausrat, T. J., Janke, C., & Kneussel, M. (2009). Synaptic activation modifies microtubules underlying transport of postsynaptic cargo. Proc Natl Acad Sci U S A, 106(21), 8731–8736. 10.1073/pnas.0812391106

52. Maeder, C. I., San-Miguel, A., Wu, E. Y., Lu, H., & Shen, K. (2014). In vivo neuron-wide analysis of synaptic vesicle precursor trafficking. Traffic, 15(3), 273–291. 10.1111/tra.12142

53. Mahmood, T., & Yang, P. C. (2012). Western blot: technique, theory, and trouble shooting. N Am J Med Sci, 4(9), 429–434. 10.4103/1947-2714.100998

54. Mathews, E. A., Stroud, D., Mullen, G. P., Gavriilidis, G., Duerr, J. S., Rand, J. B., & Hodgkin, J. (2021). Allele-specific suppression in Caenorhabditis elegans reveals details of EMS mutagenesis and a possible moonlighting interaction between the vesicular acetylcholine transporter and ERD2 receptors. Genetics, 218(4). 10.1093/genetics/iyab065

55. Mazzetti, S., Giampietro, F., Calogero, A. M., Isilgan, H. B., Gagliardi, G., Rolando, C., Cantele, F., Ascagni, M., Bramerio, M., Giaccone, G., Isaias, I. U., Pezzoli, G., & Cappelletti, G. (2024). Linking acetylated alpha-Tubulin redistribution to alpha-Synuclein pathology in brain of Parkinson’s disease patients. NPJ Parkinsons Dis, 10(1), 2. 10.1038/s41531-023-00607-9

56. McIntire, S. L., Jorgensen, E., Kaplan, J., & Horvitz, H. R. (1993). The GABAergic nervous system of Caenorhabditis elegans. Nature, 364(6435), 337–341. 10.1038/364337a0

57. Mello, C. C., Kramer, J. M., Stinchcomb, D., & Ambros, V. (1991). Efficient gene transfer in C.elegans: extrachromosomal maintenance and integration of transforming sequences. EMBO J, 10(12), 3959–3970. http://www.ncbi.nlm.nih.gov/entrez/query.fcgi?cmd=Retrieve&db=PubMed&dopt=Citation&list_uids=1935914

58. Millecamps, S., & Julien, J. P. (2013). Axonal transport deficits and neurodegenerative diseases. Nat Rev Neurosci, 14(3), 161–176. 10.1038/nrn3380

59. Mondal, S., Ahlawat, S., Rau, K., Venkataraman, V., & Koushika, S. P. (2011). Imaging in vivo neuronal transport in genetic model organisms using microfluidic devices. Traffic, 12(4), 372–385. 10.1111/j.1600-0854.2010.01157.x

60. Muniesh, M. S., Barmaver, S. N., Huang, H. Y., Bayansan, O., & Wagner, O. I. (2020). PTP-3 phosphatase promotes intramolecular folding of SYD-2 to inactivate kinesin-3 UNC-104 in neurons. Mol Biol Cell, 31(26), 2932–2947. 10.1091/mbc.E19-10-0591

61. Nass, R., Hall, D. H., Miller, D. M., 3rd, & Blakely, R. D. (2002). Neurotoxin-induced degeneration of dopamine neurons in Caenorhabditis elegans. Proc Natl Acad Sci U S A, 99(5), 3264–3269. 10.1073/pnas.042497999

62. Niwa, S., Lipton, D. M., Morikawa, M., Zhao, C., Hirokawa, N., Lu, H., & Shen, K. (2016). Autoinhibition of a Neuronal Kinesin UNC-104/KIF1A Regulates the Size and Density of Synapses. Cell Rep, 16(8), 2129–2141. 10.1016/j.celrep.2016.07.043

63. O’Hagan, R., Avrutis, A., & Ramicevic, E. (2022). Functions of the tubulin code in the C. elegans nervous system. Mol Cell Neurosci, 123, 103790. 10.1016/j.mcn.2022.103790

64. Pereira, L., Kratsios, P., Serrano-Saiz, E., Sheftel, H., Mayo, A. E., Hall, D. H., White, J. G., LeBoeuf, B., Garcia, L. R., Alon, U., & Hobert, O. (2015). A cellular and regulatory map of the cholinergic nervous system of C. elegans. Elife, 4. 10.7554/eLife.12432

65. Peris, L., Wagenbach, M., Lafanechere, L., Brocard, J., Moore, A. T., Kozielski, F., Job, D., Wordeman, L., & Andrieux, A. (2009). Motor-dependent microtubule disassembly driven by tubulin tyrosination. J Cell Biol, 185(7), 1159–1166. 10.1083/jcb.200902142

66. Reed, N. A., Cai, D., Blasius, T. L., Jih, G. T., Meyhofer, E., Gaertig, J., & Verhey, K. J. (2006). Microtubule acetylation promotes kinesin-1 binding and transport. Curr Biol, 16(21), 2166–2172. http://www.ncbi.nlm.nih.gov/entrez/query.fcgi?cmd=Retrieve&db=PubMed&dopt=Citation&list_uids=17084703

67. Sandoval, G. M., Duerr, J. S., Hodgkin, J., Rand, J. B., & Ruvkun, G. (2006). A genetic interaction between the vesicular acetylcholine transporter VAChT/UNC-17 and synaptobrevin/SNB-1 in C. elegans. Nat Neurosci, 9(5), 599–601. 10.1038/nn1685

68. Shyu, Y. J., Hiatt, S. M., Duren, H. M., Ellis, R. E., Kerppola, T. K., & Hu, C. D. (2008). Visualization of protein interactions in living Caenorhabditis elegans using bimolecular fluorescence complementation analysis. Nat Protoc, 3(4), 588–596. https://doi.org/nprot.2008.16 [pii] 10.1038/nprot.2008.16

69. Smith, R., Chung, H., Rundquist, S., Maat-Schieman, M. L., Colgan, L., Englund, E., Liu, Y. J., Roos, R. A., Faull, R. L., Brundin, P., & Li, J. Y. (2006). Cholinergic neuronal defect without cell loss in Huntington’s disease. Hum Mol Genet, 15(21), 3119–3131. 10.1093/hmg/ddl252

70. Solinger, J. A., Paolinelli, R., Kloss, H., Scorza, F. B., Marchesi, S., Sauder, U., Mitsushima, D., Capuani, F., Sturzenbaum, S. R., & Cassata, G. (2010). The Caenorhabditis elegans Elongator complex regulates neuronal alpha-tubulin acetylation. PLoS Genet, 6(1), e1000820. 10.1371/journal.pgen.1000820

71. Surana, S., Villarroel-Campos, D., Lazo, O. M., Moretto, E., Tosolini, A. P., Rhymes, E. R., Richter, S., Sleigh, J. N., & Schiavo, G. (2020). The evolution of the axonal transport toolkit. Traffic, 21(1), 13–33. 10.1111/tra.12710

72. Suzuki, M., Desmond, T. J., Albin, R. L., & Frey, K. A. (2001). Vesicular neurotransmitter transporters in Huntington’s disease: initial observations and comparison with traditional synaptic markers. Synapse, 41(4), 329–336. 10.1002/syn.1089

73. Sze, J. Y., Victor, M., Loer, C., Shi, Y., & Ruvkun, G. (2000). Food and metabolic signalling defects in a Caenorhabditis elegans serotonin-synthesis mutant. Nature, 403(6769), 560–564. 10.1038/35000609

74. Teoh, J. S., Vasudevan, A., Wang, W., Dhananjay, S., Chandhok, G., Pocock, R., Koushika, S. P., & Neumann, B. (2022). Synaptic branch stability is mediated by non-enzymatic functions of MEC-17/alphaTAT1 and ATAT-2. Sci Rep, 12(1), 14003. 10.1038/s41598-022-18333-2

75. Tien, N. W., Wu, G. H., Hsu, C. C., Chang, C. Y., & Wagner, O. I. (2011). Tau/PTL-1 associates with kinesin-3 KIF1A/UNC-104 and affects the motor’s motility characteristics in C. elegans neurons. Neurobiol Dis, 43(2), 495–506. 10.1016/j.nbd.2011.04.023

76. Wagner, O. I., Esposito, A., Kohler, B., Chen, C. W., Shen, C. P., Wu, G. H., Butkevich, E., Mandalapu, S., Wenzel, D., Wouters, F. S., & Klopfenstein, D. R. (2009). Synaptic scaffolding protein SYD-2 clusters and activates kinesin-3 UNC-104 in C. elegans. Proc Natl Acad Sci U S A, 106(46), 19605–19610. 10.1073/pnas.0902949106

77. Walter, W. J., Beránek, V., Fischermeier, E., & Diez, S. (2012). Tubulin acetylation alone does not affect kinesin-1 velocity and run length in vitro. PLoS One, 7(8), e42218. 10.1371/journal.pone.0042218

78. Wloga, D., Joachimiak, E., & Fabczak, H. (2017). Tubulin Post-Translational Modifications and Microtubule Dynamics. Int J Mol Sci, 18(10). 10.3390/ijms18102207

79. Wu, G. H., Muthaiyan Shanmugam, M., Bhan, P., Huang, Y. H., & Wagner, O. I. (2016). Identification and Characterization of LIN-2(CASK) as a Regulator of Kinesin-3 UNC-104(KIF1A) Motility and Clustering in Neurons. Traffic, 17(8), 891–907. 10.1111/tra.12413

80. Xu, F., Takahashi, H., Tanaka, Y., Ichinose, S., Niwa, S., Wicklund, M. P., & Hirokawa, N. (2018). KIF1Bbeta mutations detected in hereditary neuropathy impair IGF1R transport and axon growth. J Cell Biol, 217(10), 3480–3496. 10.1083/jcb.201801085

81. Yonekawa, Y., Harada, A., Okada, Y., Funakoshi, T., Kanai, Y., Takei, Y., Terada, S., Noda, T., & Hirokawa, N. (1998). Defect in synaptic vesicle precursor transport and neuronal cell death in KIF1A motor protein-deficient mice. J Cell Biol, 141(2), 431–441. http://www.ncbi.nlm.nih.gov/entrez/query.fcgi?cmd=Retrieve&db=PubMed&dopt=Citation&list_uids=9548721

82. Zadrozny, M., Drapich, P., Gasiorowska-Bien, A., Niewiadomski, W., Harrington, C. R., Wischik, C. M., Riedel, G., & Niewiadomska, G. (2024). Neuroprotection of Cholinergic Neurons with a Tau Aggregation Inhibitor and Rivastigmine in an Alzheimer’s-like Tauopathy Mouse Model. Cells, 13(7). 10.3390/cells13070642

83. Zhang, Y. V., Hannan, S. B., Stapper, Z. A., Kern, J. V., Jahn, T. R., & Rasse, T. M. (2016). The Drosophila KIF1A Homolog unc-104 Is Important for Site-Specific Synapse Maturation. Front Cell Neurosci, 10, 207. 10.3389/fncel.2016.00207

84. Zhu, H., Duerr, J. S., Varoqui, H., McManus, J. R., Rand, J. B., & Erickson, J. D. (2001). Analysis of point mutants in the Caenorhabditis elegans vesicular acetylcholine transporter reveals domains involved in substrate translocation. J Biol Chem, 276(45), 41580–41587. 10.1074/jbc.M103550200

